# Zinc Dependent Conformational Changes in the Cation Diffusion Facilitator YiiP from S. oneidensis

**DOI:** 10.1101/2020.12.29.424758

**Authors:** Maria Lopez-Redondo, Shujie Fan, Akiko Koide, Shohei Koide, Oliver Beckstein, David L. Stokes

## Abstract

YiiP is a secondary transporter that couples Zn^2+^ transport to the proton motive force. Structural studies of YiiP from prokaryotes as well as Znt8 from humans revealed three different Zn^2+^ sites and a conserved homodimeric architecture. These structures define the inward-facing and outward-facing states that characterize the archetypal alternating access mechanism of transport. To study effects of Zn^2+^ binding on the conformational transition, we have used a YiiP/Fab complex for single-particle cryo-EM together with Molecular Dynamics simulation to compare structures of YiiP from *S. oneidensis* in presence and absence of Zn^2+^. Without Zn^2+^, YiiP exhibits enhanced flexibility and adopts a novel conformation that appears to be an intermediate state. The transition closes a hydrophobic gate and is controlled by the Zn^2+^ site at the cytoplasmic membrane interface. This work enhances our understanding of individual Zn^2+^ binding sites and their role in the conformational dynamics that governs the transport cycle.

## Introduction

YiiP is a Zn^2+^ transporter from bacteria belonging to the family of Cation Diffusion Facilitators (CDF), also known as SLC30. YiiP functions as a Zn^2+^/H^+^ antiporter and is capable of using the proton motive force to remove Zn^2+^ from the cytoplasm (Cotrim et al., 2019). Related members of the CDF family are present in all kingdoms of life; although Zn^2+^ is a prevalent substrate, Mn^2+^, Co^2+^, Fe^2+^, Ni^2+^ and Cd^2+^ are other transition metal ions associated with this family (Cubillas et al., 2013; Montanini et al., 2007). Human homologs include a group of ten transporters named Znt1-10 which are largely responsible for loading various organelles with Zn^2+^ (Kambe, 2012). There is increasing appreciation for the physiological roles played by Zn^2+^ in a number of biological processes, including synaptic transmission, oocyte fertilization and insulin secretion (Liang et al., 2016). It has been estimated that 10% of the mammalian proteome is bound to Zn^2+^, either as a co-factor for enzymatic reactions or as a structural element stabilizing a protein fold (Andreini et al., 2006). Insufficient Zn^2+^ in the diet has significant health consequences, including chronic inflammation, diarrhea, and growth retardation (Prasad, 2013) and genetic defects in transporters have been associated with diabetes and Alzheimers disease (Lovell et al., 2005; Sladek et al., 2007). Despite relatively high (millimolar) total concentrations of Zn in the cell, levels of free Zn^2+^ are in the picomolar range (Maret, 2013), due to the high binding capacity of the proteome together with the inducible expression of metallothionein (Kimura and Kambe, 2016). Given these unique physiological considerations, a coordinated network of transporters, which also include ZIP family transporters (SLC39A), P-type ATPase (ZntA) and ABC transporters (Cotrim et al., 2019; Neupane et al., 2019) is required to maintain homeostasis of the organism.

CDF transporters share a common architecture, which is presumed to underlie a common mechanism of transport. X-ray structures of YiiP from *E. coli* (ecYiiP) (Lu et al., 2009; Lu and Fu, 2007) revealed a homodimeric assembly in which each protomer is divided into a transmembrane domain (TMD) composed of six helices and a C-terminal cytoplasmic domain (CTD) with a fold resembling a metallochaperone. Conserved metal ion binding sites are seen in the TMD (site A), which is thought to function as the transport site, and in the CTD (site C). YiiP also has a non-conserved site in the M2-M3 intracellular loop (site B) as well as a second metal ion bound by non-conserved residues at site C, making a total of four Zn^2+^ ions bound per protomer. Although other CDF family members conform to this overall architecture, many include a His-rich intracellular loop between M4 and M5 that has been postulated to play a role in delivering Zn^2+^ to the transport sites (Podar et al., 2012). Several X-ray structures have been determined of the truncated CTD from various species, which show that it dimerizes on its own and undergoes a scissor-like conformational change upon binding Zn^2+^ (Cotrim et al., 2019). Cryo-EM structures of the intact YiiP dimer from *S. oneidensis* (soYiiP) were determined from ordered arrays in a lipid membrane, which confirmed the overall architecture of the *E. coli* protein but revealed major conformational changes. Specifically, accessibility of transport sites indicated that the X-ray structure was in an outward-facing state, whereas the cryo-EM structure was an inward-facing state. In addition, the comparison highlighted a large scissor-like displacement of the TMD from each protomer that was initially implicated in transport. However, di-sulfide crosslinks engineered to prevent these movements had no effect on transport function, suggesting that more subtle conformational changes may be at play (Lopez-Redondo et al., 2018). Hydroxyl radical labeling has been used to identify hydrophobic residues at the cytoplasmic side of M5 and M6 that were proposed to gate access to the transport sites (site A), implying a rocking of the outer membrane helices (M1,2,4,5) relative to the inner membrane helices (M3,6) mediating the dimer interface (Gupta et al., 2014). Recent Molecular Dynamic (MD) simulations using a composite model for YiiP and cryo-EM structures of the mammalian transporter Znt8 provide additional support for these motions and for the ability of the hydrophobic gate to control access (Sala et al., 2019; Xue et al., 2020). In addition, the cryo-EM structure revealed two unexpected Zn binding sites; although not conserved based on amino acid sequence, the two new sites appear in similar locations to site B and site C on YiiP and the former was proposed to play an active role in recruiting ions for transport.

For the current work, we have explored the effects of Zn^2+^ binding on the conformational dynamics of soYiiP using cryo-EM and MD simulation. To facilitate the cryo-EM analysis, we used a phage display screen to develop an antibody Fab fragment recognizing the CTD. After using an *in vitro* transport assay to verify that the Fab did not interfere with function, we generated several cryo-EM structures before and after chelation of Zn^2+^ ions. 3D classification and 3D variability analysis of cryo-EM datasets revealed conformational variability within individual samples, which appear to correlate with Zn^2+^ binding to the site on the TM2-TM3 loop (site B). Two discrete structures revealed distinct conformations corresponding to holo and apo states, whereas 3D variability analysis elucidated the transition between these states. The refined structures, at 3.4 and 4.0 Å resolution, respectively, revealed a disordering of the TM2-TM3 loop, as well as Zn site B, and a closing of the hydrophobic gate when Zn^2+^ was removed. For MD simulation, we developed a non-bonded dummy model for Zn^2+^ ions and validated it against known Zn^2+^-binding proteins. MD simulations of the apo and holo states of YiiP revealed enhanced dynamics in the absence of Zn^2+^ that we correlated with the cryo-EM structures. In particular, enhanced movements were observed for the M2/M3 loop and for the CTD domain relative to the TM domain in the absence of Zn^2+^. Despite the enhanced flexibility and gate closure, dimer interfaces within the TMD and the CTD were preserved in both cryo-EM structures and the MD simulations.

## Results

### Zn^2+^ is required for formation of tubular crystals

In previous work, we employed an automated screening pipeline to produce tubular crystals of YiiP within reconstituted bilayers (Kim et al., 2010). Cryo-EM and helical reconstruction were then used to generate structures of the two-fold symmetric homodimer in a membrane environment (Coudray et al., 2013; Lopez-Redondo et al., 2018). Although Zn^2+^ was not added during purification or crystallization, a high resolution structure revealed strong densities at each of the four metal ion binding sites (Lopez-Redondo et al., 2018), suggesting that metal ions either co-purified with the protein or were scavenged from buffers used for purification and crystallization. As a first attempt to evaluate the structural effects of removing metal ions, we treated samples with 5 mM EDTA for 10 min prior to reconstitution and crystallization (Table I). After EDTA treatment, tubular crystals readily formed under standard conditions, which involved two weeks of dialysis in buffers lacking both Zn^2+^ and EDTA (Suppl. Fig. 1). After imaging these samples by cryo-EM, 2D classification revealed the prevalence of two helical morphologies characterized by D1 and D3 symmetries, as has been previously described (Lopez-Redondo et al., 2018). The global resolution of these structures (referred to as 2DX) was 4.2 Å resolution though, as previously observed, the local resolution within the TMD was considerably better than for the CTD. In order to evaluate the status of the metal binding sites in these structures, the previous model from tubular crystals (PDBID 5VRF) was docked as a rigid body to these new density maps, thus illustrating that the conformation was unaffected by the EDTA treatment (Fig. 1). In addition, densities consistent with a bound ion and with coordinating side chains were clearly visible at Zn sites A and B; the lower resolution of the CTD made site C difficult to evaluate. This result suggests either that the off-rate for metal ions is extremely low or that the affinity is high enough to scavenge trace metal ions from the dialysis buffers used to produce the tubular crystals. We then attempted to produce the Zn^2+^-free (apo) state by including 0.5 mM EDTA in the dialysis buffer used to produce the tubular crystals. However, ordered arrays did not form under these conditions, giving the first indication that effective removal of Zn^2+^ may give rise to a conformational change that is large enough to preclude crystallization.

**Figure 1.**
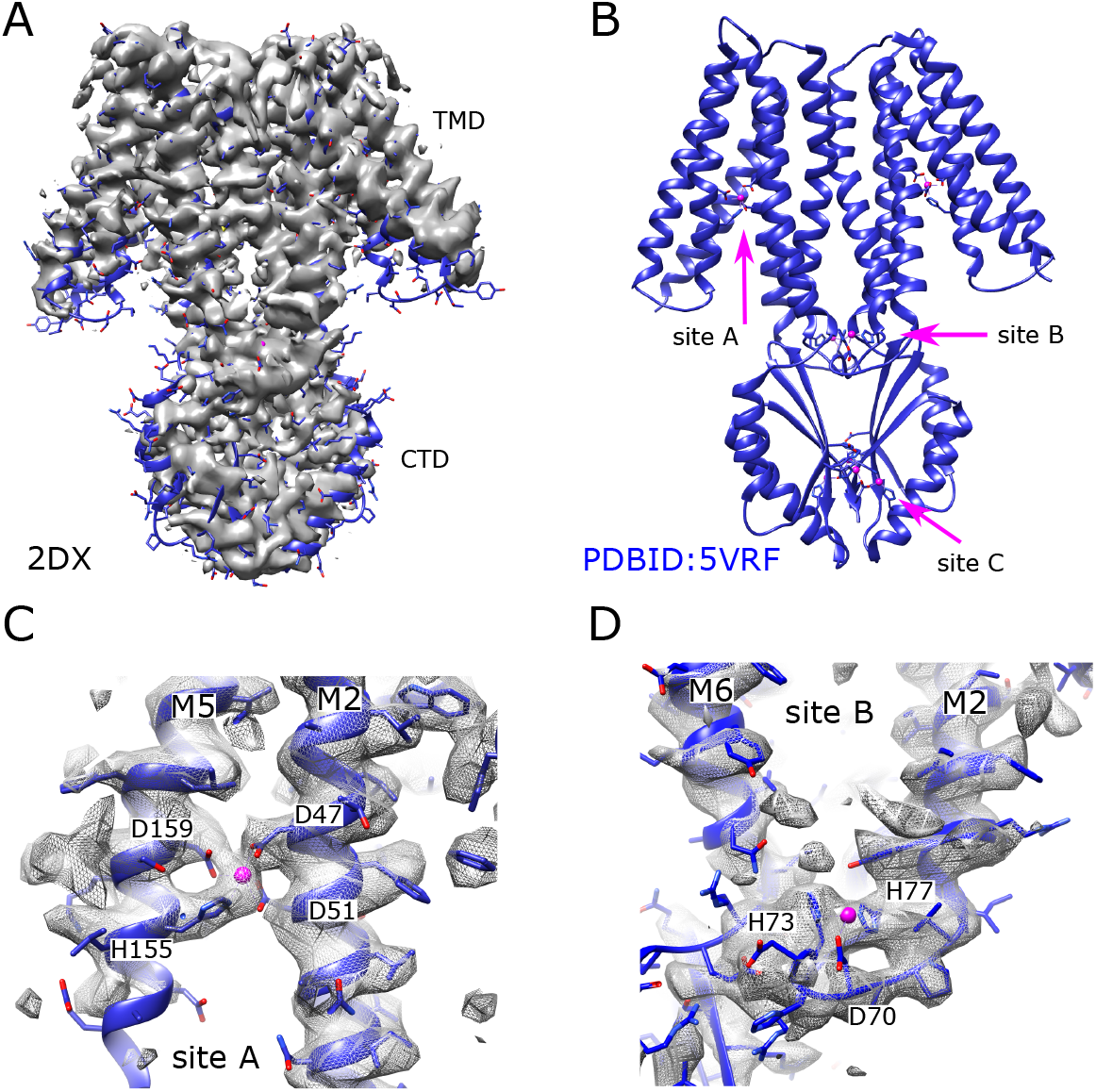
Cryo-EM structure (2DX) of EDTA-treated YiiP from tubular crystals. (A) Density map at 4.2 Å resolution of an isolated YiiP dimer masked from 3D reconstruction of tubular crystals from EDTA-treated wild-type YiiP. The previous structure from untreated tubular crystals (PDBID 5VRF) was docked as a rigid body, illustrating the similarity in the conformation. (B) The previous structure (5VRF) derived from tubular crystals showing the overall architecture of the dimer and the location of the three Zn binding sites. Zn ions are shown as pink spheres. (C) Detailed view of Zn binding site A at which the map from EDTA-treated YiiP reveals strong density, indicating that a metal ion is bound at this site. (D) Detailed view of Zn binding site B showing an ordered loop between M2 and M3 and density associated with the metal ion and coordinating residues. Side chains do not perfectly match the density, because 5VRF was only fitted as a rigid body and not explicitly refined to the map.

**Table 1.** Samples used for cryo-EM analysis

| Sample | Complex | Pre-treatment | SEC | Grid prep | Images/Particles | Res, Sym, Structure |
| --- | --- | --- | --- | --- | --- | --- |
| 2DX | YiiP helical crystals | 5mM EDTA<br>10 min | No EDTA | dialysis/<br>reconstitution |  | 4.2 Å, D3<br>4.2 Å D1,<br>2DX |
| SP1 | YiiP+Fab2 | No EDTA | No EDTA | Conc | 4312<br>102,390 | 3.8 Å, C2,<br>SP1 |
| SP2 | YiiP+Fab2r | 0.5 mM<br>EDTA o/n<sup>1</sup> | 0.5 mM<br>EDTA | Conc | 1945<br>140,465 | 4.0 Å, C1,<br>SP2 |
| SP3 | YiiP+Fab2r | 0.5 mM<br>EDTA o/n | No EDTA | Conc, Filter | 2898<br>151,898<br>85,207 | 3.4 Å, C2,<br>SP3sym<br>4.2 Å C1,<br>SP3bent |
| SP4 | YiiP+Fab2r | 0.5 mM<br>EDTA<br>0.5 mM<br>TPEN o/n | 0.5 mM<br>EDTA 0.5<br>mM TPEN | Conc, Filter | 8234<br>287,608<br>140,971 | 3.5 Å, C2,<br>SP4sym<br>4.7 Å, C1,<br>SP4bent |

<sup>1</sup> o/n indicates overnight incubation

### Development of an antibody fragment for cryo-EM analysis

Single particle cryo-EM is an alternative approach for evaluating the structure of YiiP, in which buffer conditions can be manipulated without the physical or chemical constraints of crystallization. However, the 65 kD mass for the YiiP dimer makes it challenging to determine a cryo-EM structure at near-atomic resolution. We therefore screened a phage-display library to identify Fab clones that bind to YiiP with high affinity, thus increasing the size of the complex. The phage-display library, which contains ~10^11^ unique Fab fragments (Miller et al., 2012; Sauer et al., 2020), was screened using purified YiiP reconstituted into biotinylated nanodiscs (Dominik and Kossiakoff, 2015). The library sorting process included a negative selection against empty nanodiscs to exclude nanodisc binders. The various candidate Fab’s were initially evaluated by ELISA (Fig. 2a) and those demonstrating specific binding to YiiP were sequenced and cloned into a vector for expression in *E. coli*. In this way, we identified four Fab clones that were purified in large quantities suitable for biophysical evaluation. Specifically, formation of a stable complex was examined using size-exclusion chromatography (SEC), where a shift in the elution volume indicated the presence of a stable complex between YiiP and the Fab fragment (Fig. 2b). Fab2 showed a single, uniform SEC peak corresponding to the complex and was therefore was selected for our cryo-EM work. Although initial complexes were formed using the 4D5 Fab framework employed for the phage-display library (Miller et al., 2012), we later introduced mutations into the hinge between variable and constant domains in the heavy chain (VH and CH) in an attempt to increase rigidity and thus the resolution of cryo-EM structures (Bailey et al., 2018). In particular, we substituted the SSAST sequence in the heavy chain with FNQI to generate a construct denoted Fab2r. SEC profiles and cryo-EM analysis discussed below indicated that this substitution had no negative effect on complex formation with YiiP.

**Figure 2.**
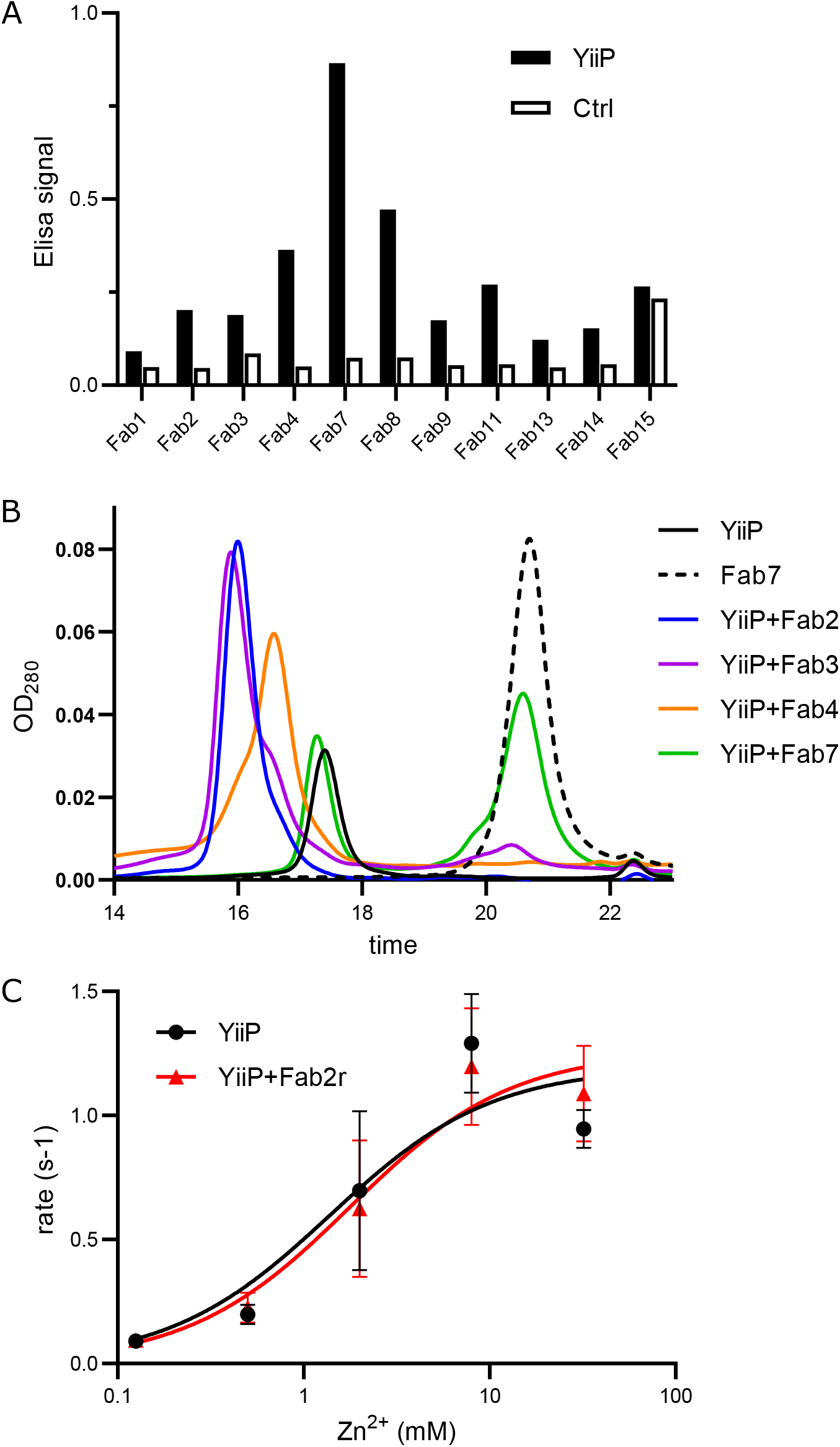
Development of antibody fragments that recognize YiiP. (A) Screening was based on phage display of a large library of Fab constructs against YiiP reconstituted in nanodiscs. Results from ELISA assay for several candidates show preferential binding to YiiP embedded in nanodiscs compared with empty nanodiscs, labeled as control. (B) Formation of a stable complex between detergent-solubilized YiiP and purified Fab candidates was evaluated by SEC. A distinct shift in elution time for Fab2 and Fab3 indicated the presence of a stable complex, whereas Fab7 did not produce a complex at all. Fab4 produced smaller shift with an asymmetric peak, probably indicating non-stoichiometric binding of Fab’s to the YiiP dimer. (C) Transport assays in presence and absence Fab2r revealed that complex formation does not affect transport activity of WT YiiP. Fitting of the data (recorded in triplicate) with the Michaelis Menton equation yielded K_M_ = 1.4 and 1.8 mM, V_max_ = 1.2 and 1.3 sec^−1^ for YiiP and the YiiP/Fab2R complex, respectively. These differences were well within the 95% confidence interval and therefore not statistically significant.

Prior to structural studies, a transport assay was used to ensure that Fab binding did not interfere with the functionality of YiiP, and hence its ability to undergo conformational changes associated with transport. For this assay, the YiiP/Fab2r complex was formed by incubation at a 2:1 molar ratio followed by reconstitution into proteoliposomes using established procedures (Lopez-Redondo et al., 2018). The results (Fig. 2c) show that the Fab had no apparent effect on either K_M_ or V_max_, suggesting that Fab binding did not interfere with conformational changes associated with the transport process.

### Development of a Zn model for MD simulation

To complement cryo-EM studies, we used MD simulation to assess the effect of Zn^2+^ on the structural dynamics of YiiP. To start, we developed a representation of metal ions that freely exchange between binding sites and the aqueous solvent, which has proved challenging for classical force fields (Li and Merz, 2017). Similar to previous work (Duarte et al., 2014), we parametrized a non-bonded dummy atom model for Zn(II) ions for the CHARMM force field and validated its performance in simulations with a selection of well-characterized proteins. Our dummy model consisted of a central Zn atom surrounded by six dummy atoms forming an octahedral shell (Suppl. Fig. 2a). The van der Waals parameters of the atoms were optimized using a genetic algorithm (Eiben et al., 1994) to reproduce a hydration energy of Zn^2+^ ions in water of −1955 kJ/mol (Marcus, 1991) and an ion-water oxygen distance of 2.08 Å (Marcus, 1988). After six generations, the optimization converged to within 1% and 0.3% of the target values, respectively (Suppl. Fig. 2b); the model also exhibited the experimentally observed coordination of six oxygen atoms in water (Marcus, 1988) (Suppl. Fig. 2c). The model was then validated by assessing how well it reproduced binding site coordination in experimental crystal structures. We performed 200 ns simulations (three repeats each) using the X-ray crystal structures for β-1,3-1,4-endoglucanase (PDBID 1U0A), stromelysin-1 (2USN) and the δ’ subunit of *E. coli* clamp-loader complex (1A5T). These structures were chosen for their high resolution, 1.6, 2.2 and 2.2 Å respectively, and for the diversity in their coordinating residues, namely HHDD, HHHD, and CCCC. His (H) and Asp (D) residues are most relevant given their presence in the binding sites of YiiP; the Cys (C) site was included for completeness. For the simulations, the Zn ion was placed at the site according to the X-ray structures followed by all-atom simulations in water. In all three cases, the Zn ion remained stably bound at the binding site, despite movements of protein elements around the site. Radial distribution functions for the Zn ion bound by His and Asp residues showed that distances to coordinating oxygen or nitrogen atoms were within 0.1 Å and 0.2 Å of the X-ray structure, respectively, indicating that the dummy model was highly suitable for these sites (Suppl. Fig. 2d-i). On the other hand, the Zn ion bound by Cys residues displayed slightly larger distance disparities (0.25-0.3 Å, Suppl. Fig. 2j-l). The excellent performance for O is expected due to reliance on Zn-oxygen interactions for the parametrization process and the very good performance for N is probably due to the overall good balance of the force field. On the other hand, larger differences for S atoms indicate that further improvements for sulfur-based sites may be warranted, for example by including sulfur compounds in the parametrization process. Nevertheless, the exclusive presence of His and Asp residues in the Zn^2+^ binding sites of YiiP means that the current version of our CHARMM Zn non-bonded dummy model is well suited for our simulations.

### Structure of the YiiP/Fab complex in the holo state

Single-particle cryo-EM was used to generate a structure of the YiiP/Fab2 complex at 3.8 Å resolution (Suppl. Fig. 3). This complex was produced by incubating Fab2 and YiiP at a 1:1 weight ratio for 1 hr at 20°C and then purifying the intact complex by SEC. As with the original tubular crystals, neither Zn^2+^ nor EDTA were included during the preparation of this sample (Table I). Peak fractions from SEC were concentrated and used immediately to produce grids for cryo-EM imaging, which resulted in 4312 images. Templates for particle picking were derived from a manual selection of ~1000 particles; automated picking resulted in an initial set of ~500,000 particles. Good particles were selected through 2D classification and ab initio jobs in cryoSPARC (Punjani et al., 2017), which then produced a structure with 3.94 Å resolution from a refinement job with ~160,000 particles. A heterogeneous refinement job was used to look for heterogeneity in this data set. Although the two resulting structures appeared to represent the same conformation, this job made a final selection of ~100,000 particles that produced the highest resolution structure (referred to as SP1, Table I). Whereas initial steps of classification and heterogeneous refinement were performed without symmetry, application of C2 symmetry during the final refinement improved the resolution of the final structure from 4.05 to 3.83 Å, indicating that the two-fold symmetry observed in both the X-ray structure and the cryo-EM structure from tubular crystals was preserved in these isolated particles.

The resulting structure reveals a homodimer of YiiP with Fab molecules, composed of a heavy and a light chain, bound to each CTD (Fig. 3). To build an atomic model, the published model from tubular crystals (PDBID 5VRF) was docked into the structure together with a homology model for the Fab. This initial structure was used for Molecular Dynamics Flexible Fitting (MDFF) (Trabuco et al., 2008), as implemented by Namdinator (Kidmose et al., 2019). The MDFF structure was improved by several rounds of PHENIX real-space refinement (Adams et al., 2010) and manual building in COOT (Emsley et al., 2010) (Table II). The resulting structure shows a long complementarity-determining region 3 (CDR3) of the Fab heavy chain making extensive contact with the C-terminus of YiiP. CDR1 and CDR2 of the heavy chain as well as CDR3 from the light chain interact with the first helix in the CTD of YiiP as well as the succeeding loop (residues 219-231 of YiiP). These interactions are all at the periphery of the CTD and do not appear to interfere with Zn^2+^ binding to site C, which is at the dimeric interface of the CTD’s. The resolution is higher in this region of the map (3.5 Å), suggesting that interactions between Fab’s and the CTD’s have a stabilizing influence (Suppl. Fig. 3d). As expected, the distal, constant domains of the Fab’s (CH and CL) have lower resolution (4.5 Å), presumably due to flexibility in the hinges between the variable and constant domains. Within the TMD of YiiP, the dimer interface mediated by M3 and M6 is reasonably well ordered (3.9 Å), but the bundle of peripheral helices (M1, M2, M4 and M5) is more flexible, making side chains difficult to identify in this region. Accordingly, the distribution of resolution from single particles differs markedly from tubular crystals (Suppl. Fig. 1d), which is likely influenced by crystal packing interactions and physical constraints of the membrane environment.

**Figure 3.**
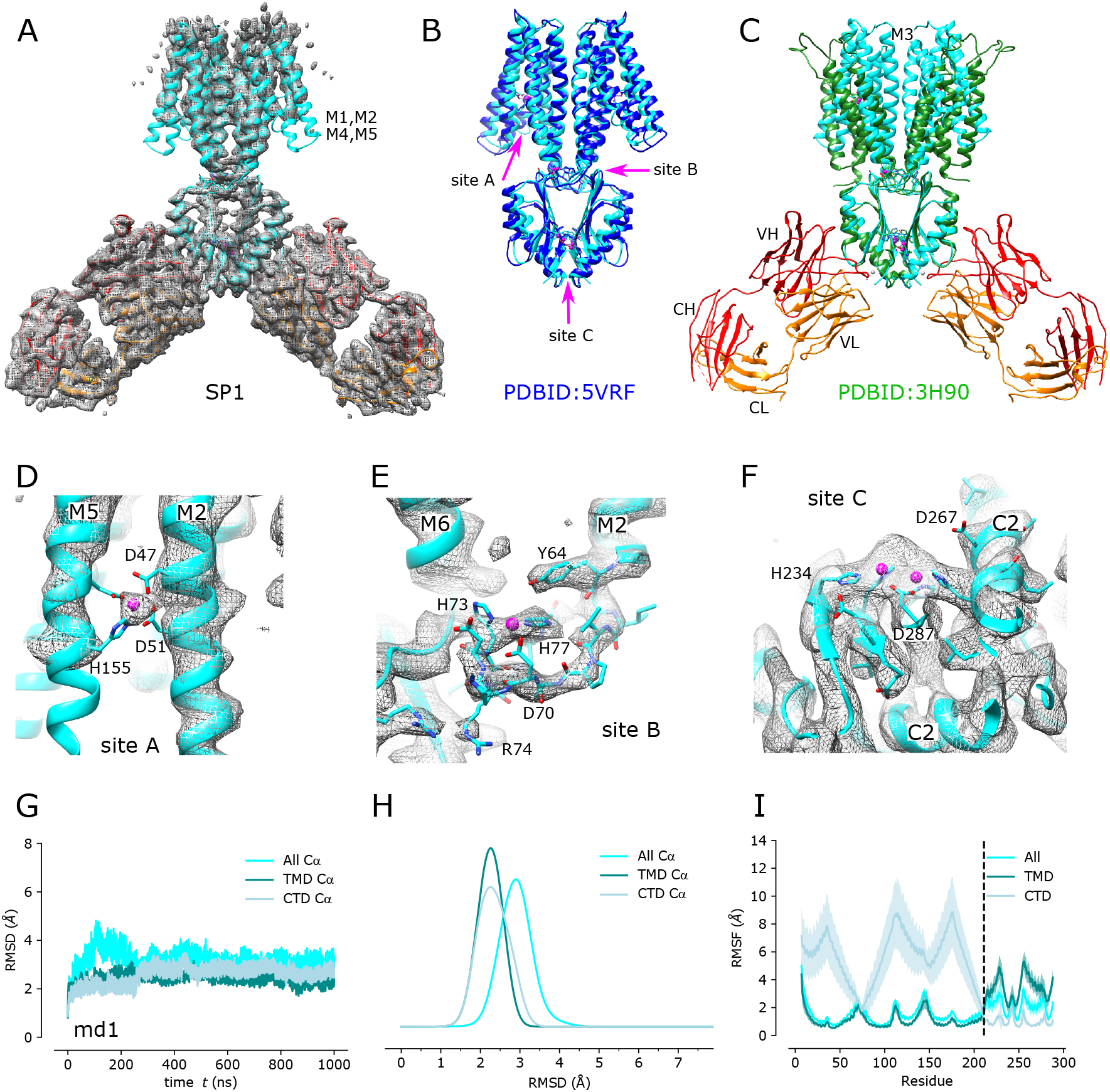
Cryo-EM structure of the untreated YiiP/Fab complex (SP1). (A) Density map at 3.8 Å overlaid with a refined atomic model with YiiP in cyan and Fab molecules in orange and red, for the light chain and heavy chain, respectively. (B) Superposition of the refined atomic model from SP1 (cyan) with the previous atomic model for YiiP (PDBID 5VRF, dark blue) showing that both models represent the same inward-facing conformation (RMSD for Cα atoms is 1.5 Å). Locations of the individual Zn sites (A, B, and C) are indicated by arrows. (C) Superposition of the SP1 model (cyan, orange, red) with the X-ray model of YiiP from *E.coli* (PDBID 3H90, green). Substantial conformational differences in the TMD are reflected in the high RMSD for Cα atoms of 10.3 Å. (D-F) Close-up views of density at the individual Zn sites in the SP1 structure. Metal ions are displayed as pink spheres. (G) C_α_ RMSD plots for one of the three MD simulations (md1) of the holo state shows relatively little change in the dimer over the course of the simulation. The three traces correspond to the different alignment schemes relative to the starting model (5VRF): “All” indicates global alignment based on the entire molecule, “TMD” indicates alignment based on the TMD only, “CTD” indicates alignment based on CTD only. (H) Distributions of RMSD derived from all three simulations using the alignment schemes indicated in the legend. These RMSD distributions, as well as those shown in other figures, were generated with a kernel density estimate (KDE). (I) RMSF plotted for each residue based on the different alignment schemes. RMSF profiles were averaged over both protomers and all three simulations, with the mean shown as the solid line and the error band indicating the standard deviation over these six profiles.

**Table 2.** CryoEM data collection and model statistics.

| Data set | SP1<br>untreated | SP2<br>EDTA<br>treated | SP3sym<br>EDTA<br>treated | SP3bent<br>EDTA<br>treated | SP4sym<br>EDTA/TPEN<br>treated | SP4bent<br>EDTA/TPEN<br>treated |
| --- | --- | --- | --- | --- | --- | --- |
| **Deposition** | N/A |  |  | N/A | N/A | N/A |
| PDB |  | 7KZX | 7KZZ |  |  |  |
| EMDB |  | EMD-23092 | EMD-23093 |  |  |  |
| **Data collection and processing** |  |  |  |  |  |  |
| Magnification | 130,000 | 130,000 | 130,000 | 130,000 | 130,000 | 130,000 |
| Voltage (kV) | 300 | 300 | 300 | 300 | 300 | 300 |
| Electron exposure (e<sup>-</sup>/Å<sup>2</sup>) | 83 | 77 | 65 | 65 | 50 | 50 |
| Defocus range (μm) | 1.0-3.0 | 1.5-3.0 | 1-2.5 | 1-2.5 | 1-2.5 | 1-2.5 |
| Pixel size (Å) | 1.035 | 1.035 | 1.048 | 1.048 | 1.079 | 1.079 |
| Symmetry imposed | C2 | C1 | C2 | C1 | C2 | C1 |
| Initial particle images<br>(no.) | 473,960 | 848,576 | 1,448,191 | 1,448,191 | 1,685,559 | 1,685,559 |
| Final particle images<br>(no.) | 102,390 | 140,565 | 151,898 | 85,207 | 287,608 | 140,971 |
| Map resolution (Å) | 3.83 | 4.00 | 3.42 | 4.18 | 3.52 | 4.66 |
| FSC threshold | 0.143 | 0.143 | 0.143 | 0.143 | 0.143 | 0.143 |
| **Model Refinement** |  |  |  | MDFF<sup>1</sup> | MDFF<sup>1</sup> | MDFF<sup>1</sup> |
| Model composition |  |  |  |  |  |  |
| Non-hydrogen atoms | 10416 | 10446 | 10470 | 10446 | 10468 | 10446 |
| Protein residues | 1362 | 1361 | 1364 | 1361 | 1364 | 1361 |
| Ligands | 8 | 0 | 8 | 0 | 0 | 0 |
| RMS deviations |  |  |  |  |  |  |
| Bond lengths (Å) | 0.004 | 0.011 | 0.014 | 0.012 | 0.012 | 0.009 |
| Bond angles (°) | 0.884 | 1.103 | 2.397 | 1.275 | 1.238 | 1.119 |
| Validation |  |  |  |  |  |  |
| MolProbity score | 2.39 | 2.31 | 2.13 | 2.73 | 2.41 | 2.65 |
| Clashscore | 20.96 | 19.88 | 2.30 | 40.62 | 21.56 | 34.86 |
| Rotamer outliers (%) | 0.00 | 0.00 | 4.11 | 0.00 | 0.00 | 0.00 |
| Ramachandran plot |  |  |  |  |  |  |
| Favored (%) | 89.16 | 91.10 | 85.00 | 86.16 | 88.66 | 86.84 |
| Allowed (%) | 10.84 | 8.60 | 11.57 | 13.61 | 11.04 | 13.16 |
| Disallowed (%) | 0.00 | 0.30 | 3.43 | 0.22 | 0.30 | 0.00 |
| Model vs. Data CC (mask) | 0.78 | 0.81 | 0.78 | 0.78 | 0.81 | 0.82 |

<sup>1</sup> Models created by Namdinator, which removes Zn ligands during its fitting process

Comparison of the refined structures from the detergent-solubilized SP1 complex and membrane-bound YiiP in the tubular crystals indicate that the YiiP dimer adopts a comparable inward-facing conformation despite the differing conditions of the two samples. Rigid-body docking of the structure from tubular crystals (5VRF) matched the map densities very well with a cross-correlation of 0.82. Furthermore, comparison of 5VRF with the atomic model refined to the SP1 map generated a low RMSD for C_α_ atoms of 1.53 Å for the entire YiiP dimer (Table III). When individual monomers were compared, the RMSD decreased slightly to 1.2 Å, indicating slight flexibility in the angle between the monomers. On the other hand, this conformation differs substantially from X-ray structures of detergent-solubilized YiiP from *E. coli* (PDBID 3H90), which show a scissor-like displacement of TMD’s that disrupts the interaction between M3 helices (Fig. 3c). Although the CTD’s are virtually identical (RMSD = 0.97 Å), the TMD’s are quite different (RMSD = 10.3 Å) due not only to the scissor-like displacement but also to rocking of the four-helix bundle (M1,M2,M4,M5) to produce the outward-facing state as has been previously described (Lopez-Redondo et al., 2018).

**Table 3.** RMSD between atomic models*

|  | 3H90 | 5VRF | SP1 | SP3<br>sym | SP4<br>sym | SP2 | SP3<br>bent | SP4<br>bent |
| --- | --- | --- | --- | --- | --- | --- | --- | --- |
| 3H90 |  |  |  |  |  |  |  |  |
| 5VRF | 8.3 |  |  |  |  |  |  |  |
| SP1 | 10.3 | 1.5 |  |  |  |  |  |  |
| SP3sym | 8.7 | 1.6 | 1.2 |  |  |  |  |  |
| SP4sym | 8.6 | 1.8 | 1.4 | 1.2 |  |  |  |  |
| SP2 | 14.3 | 4.4 | 4.4 | 4.4 | 4.1 |  |  |  |
| SP3bent | 14.2 | 4.2 | 4.0 | 3.9 | 3.8 | 1.5 |  |  |
| SP4bent | 15.0 | 4.6 | 4.5 | 4.5 | 4.2 | 1.5 | 1.8 |  |

\* values in the table correspond to RMSD for all C<sub>α</sub> atoms in Å

Like the structure from tubular crystals, the SP1 complex appeared to have metal ions bound at each of the Zn^2+^ binding sites. Strong densities were visible at each of these sites, encompassing both the ion sites themselves and the coordinating residues (Fig. 3d-f). Furthermore, the local resolution was enhanced at these sites relative to the neighboring regions of the protein (Suppl. Fig. 3d). In particular, the presence of distinct density at site A within the TMD (Fig. 3d) was particularly notable, given that the lower resolution of the peripheral membrane helices generally precluded side chain densities in this region of the map. Thus, like the structure from tubular crystals, the SP1 complex appears to represent the Zn^2+^-bound, holo state, despite the lack of exogenous Zn^2+^ in the buffers used for purification and sample preparation.

To complement this experimental approach, we used MD simulations to characterize the dynamics of YiiP in the holo state. We used 5VRF as the starting model with a Zn ion (parametrized with the non-bonded dummy model) placed at each of the eight sites in the homodimer and equilibrated this structure in a membrane composed of palmitoyloleoylphosphatidylethanolamine (POPE) and palmitoyloleoylphosphatidylglycerol (POPG) lipids in a 4:1 ratio, approximating a typical *E. coli* plasma membrane (Raetz, 1986). Three independent simulations were run for 1 μs each during which Zn^2+^ remained stably bound at all of the binding sites (Suppl. Fig. 4). RMSD’s for C_α_ atoms in the TMD, the CTD, and the whole dimer were calculated for all frames relative to the starting model. Data from all three simulations (Fig. 3g and Suppl. Fig. 5) were similar with values ranging from 2-3.5 Å. As can be seen in the distribution of RMSD’s (Fig. 3h), the average displacements were consistently higher for the whole dimer relative to values for individual domains, which is indicative of inter-domain movements. Per-residue RMSF’s were calculated after structurally superimposing frames on either the whole dimer, the TMD, or the CTD (Fig. 3i) and provide more robust evidence of inter-domain movements. In particular, profiles generated after alignment of CTD’s show 2-4x higher RMSF values for the TMD’s with peaks associated with M1-M2 and M4-M5 helix pairs. Despite these inter-domain movements, the dimer interface within each of these domains remains intact, as is discussed further below.

### Structure of the YiiP/Fab complex in the apo state

Given the high avidity of YiiP for Zn^2+^ and possibly other transition metal ions, we explored various protocols for generating the Zn^2+^-free, apo state of the YiiP/Fab complex for cryo-EM. In particular, the complex was incubated in 0.5 mM EDTA or in a combination of 0.5 mM EDTA and 0.5 mM TPEN, which bind a wide range of divalent cations with high affinity and thus expected to strip any such ions from binding sites of YiiP. For this complex, we used the rigidified Fab2r. Three independent data sets were collected from samples prepared under slightly different conditions, as summarized in Table I. The most successful procedure involved incubating the complex overnight in 0.5 mM EDTA and then purifying it in this same EDTA-containing buffer. This sample (referred to as SP2) produced a reasonably homogeneous data set and a refined structure at 4.0 Å resolution (Suppl. Fig. 6, Table II). Comparison with the holo state discussed above revealed a large-scale conformational change involving an ~25° bend between the CTD of YiiP and the membrane domain (Fig. 4a vs. Fig. 4b). This bend disrupts the C2 symmetry that characterized the holo state, although the dimeric interfaces in the CTD and the TM domain remain intact and local two-fold symmetry within these domains is preserved (Fig. 4c & d). Indeed, the overall architecture of the TMD is well preserved, especially at the dimer interface where M3 and M6 can be superimposed with RMSD of 0.7 Å for C_α_. Similarly, the CTD is relatively unchanged (RMSD of 1.99 Å) and recognition by the Fab’s was not affected by chelation of metal ions. As before, the vicinity of the CTD/Fab interface generated the highest resolution in the map with side chains clearly visible in both molecules. Despite the use of the rigidified Fab2r molecule, the local resolution of the distal, constant domains (CH and CL) remained significantly lower than that of the proximal domains (VH and VL) (Suppl. Fig. 6d), suggesting that the mutation did not completely eliminate flexibility of the hinge. The resolution in the TMD was also significantly lower than the holo structure, indicating increased flexibility and ultimately representing the limiting factor in our cryo-EM analysis.

**Figure 4.**
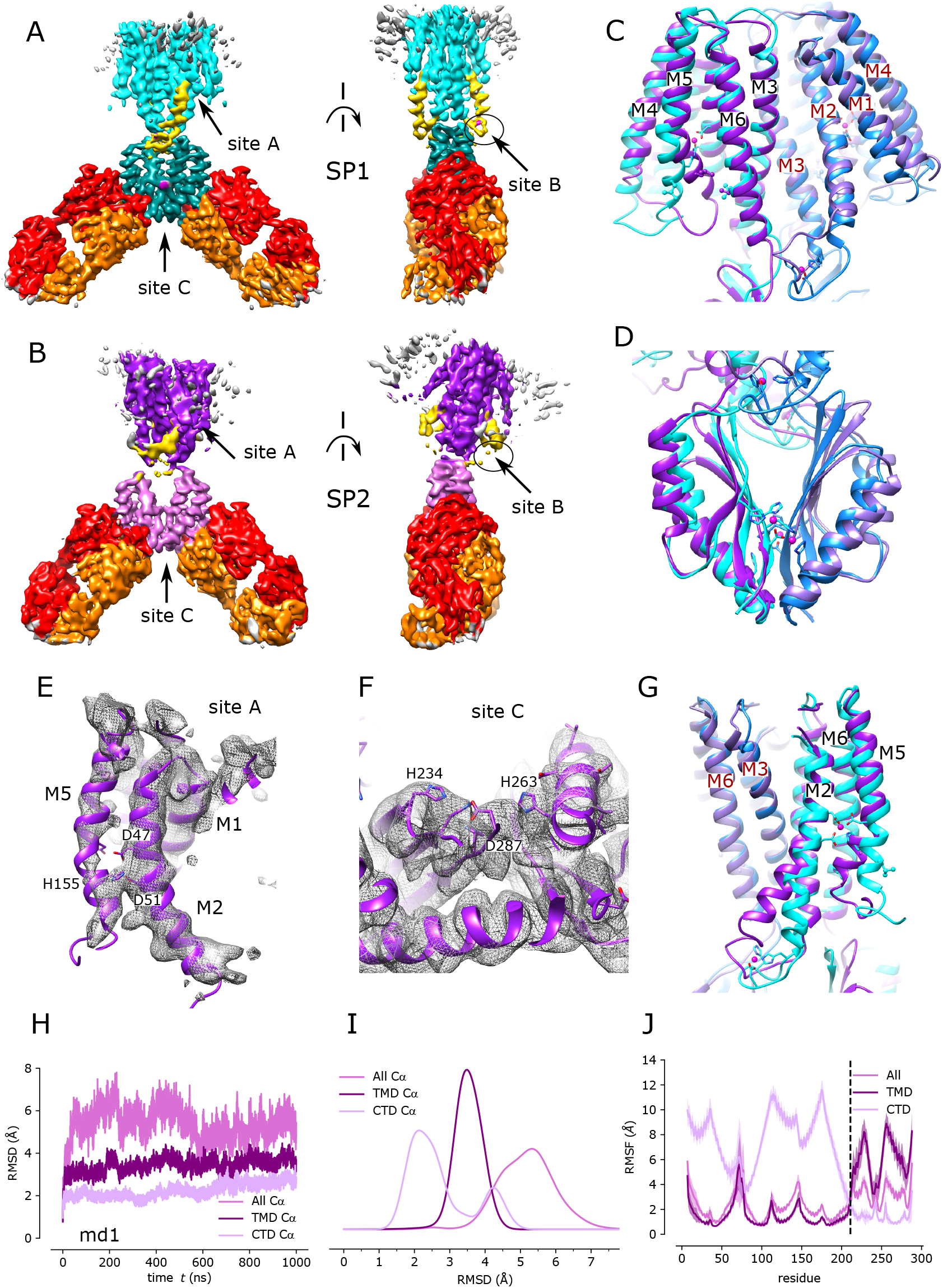
Cryo-EM structure of EDTA-treated YiiP/Fab complex (SP2) and comparison with the untreated complex (SP1). (A) SP1 structure of untreated complex showing C2 symmetry with colors highlighting the following features: light cyan for the TMD, dark cyan for the CTD, yellow for the TM2/TM3 loop, orange for the Fab light chain, red for the Fab heavy chain. The location of Zn sites A, B, and C are indicated by arrows. (B) Comparable views of the SP2 structure for the EDTA-treated complex with the following colors: dark purple for the TM domain, light purple for the CTD, yellow for the TM2/TM3 loop. (C) Alignment of TMD’s for the two structures (cyan for SP1 and purple for SP2) show preservation of the dimer interface and overall architecture of this domain. Individual protomers are shown in different shades. (D) Alignment of the CTD’s for the two structures shows preservation of the dimer interface with a modest increase in separation in the apo state. (E) Close-up of Zn site A in the TMD from the SP2 structure showing a lack of density at the ion binding site and a bend in the M2 helix near Asp51. (F) Close-up of Zn site C in the CTD with lack of density indicating an absence of ions at this site. (G) Comparison of M2 and M5 in apo and holo states shows bends originating at Zn site A. (H) C_α_ RMSD plots for one of the three MD simulations (md1) of the apo state. The three traces correspond to the different alignment schemes as indicated in the legend and described in Fig. 3. (I) Distributions of RMSD derived from all three simulations using the three alignment schemes. (J) RMSF profiles averaged over both protomers and all three simulations based on the different alignment schemes; the error bands indicate the standard deviation over these six profiles.

Despite the lower resolution, the transmembrane helices are clearly visible in the map and the overall architecture of the TMD is consistent with the inward-facing conformation seen in 2DX and SP1 structures. However, there is a significant change in the M2 helix, which undergoes a distinct bend towards its cytoplasmic end (Fig. 4e,g). This change is accompanied by a complete disordering of the M2/M3 loop, which normally carries a Zn^2+^ binding site (site B). In addition, the cytoplasmic end of M5 is bent such that it closes the gap between M6 (Fig. 4g). The bends in M2 and M5 both occur near residues that form Zn site A: Asp51 and His155. No densities are observed at Zn^2+^ binding sites within the membrane (site A) or in the CTD (site C) (Fig. 4e & f) indicating that EDTA treatment was indeed effective in stripping metal ions from these sites. The lack of density for the M2/M3 loop made it impossible to evaluate whether Zn^2+^ was bound at site B, but the disordering of coordinating residues within this loop is a very plausible result of removing the metal ions from this site. Thus, we believe that this structure represents the apo state of the YiiP.

MD simulations were used to evaluate the dynamics of the apo state. Similar to the holo state, we used 5VRF as a starting model for three independent μsec-long simulations, in which Zn ions were removed from all of the binding sites. As with the holo structure, RMSD’s and RMSF’s were lower for individual domains relative to the whole dimer (Fig. 4h-j and Suppl. Fig. 7). The RMSD values were ~2-fold higher for the apo state compared to the holo state, with the exception of the CTD, which displayed a lesser increase. In order to compare holo and apo simulations more directly, we plotted RMSF values for aligned TMD’s and aligned CTD’s for each monomer (Fig. 5a). As might be expected, these plots show distinct peaks for loops connecting secondary structure elements in both TMD and CTD. It is notable that the M2/M3 loop undergoes a dramatic increase in flexibility in the apo state, which correlates with the disordering of this loop in the cryo-EM structures. Increased flexibility in the apo simulations is also evident from the increased RMSD for C_α_ atoms in the loop, from <1 Å for the holo state to ~4 Å in the absence of a stabilizing ion (Fig. 5b and Suppl. Fig. 8). The differing mobility is depicted in Fig. 5c & d, which shows the range of conformations sampled by the M2/M3 loop in the two simulations. We also quantified stability of the dimer interface and angles between TMD and CTD during the simulations (Fig. 6). Like the cryo-EM structures, dimeric interfaces in the simulations remained largely intact, especially in the TMD where the fraction of residues maintaining native contacts is close to 1 (Fig. 6a). Contacts within the CTD are somewhat less stable and deteriorate in the apo state although the overall architecture of the dimer remains intact (Fig. 6b & d). The hinge between the TMD and CTD also has enhance flexibility in the apo state with a mean angle of ~30° compared to ~15° for the holo state (Fig. 6e).

**Figure 5.**
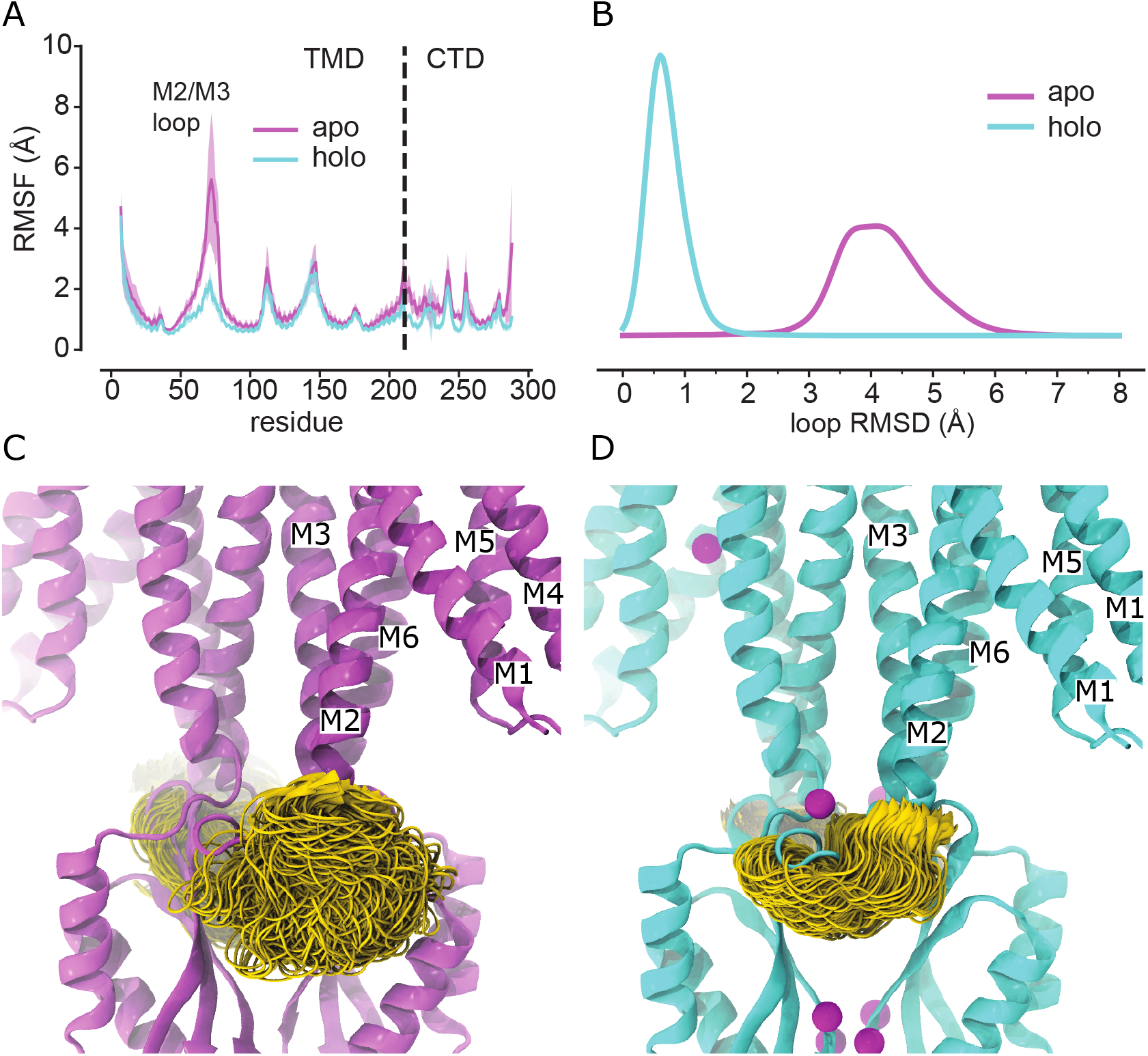
MD simulations show greater flexibility of the M2/M3 loop in the apo state. (A) RMSF calculated for each YiiP monomer for all of the MD simulations in apo state (purple) and the holo state (cyan). RMSF’s for residues in the TMD or CTD, indicated by the dashed line, was calculated after superposition on the corresponding domain. RMSF profiles were averaged over the two protomers and over the three independent simulations, with the error bands indicating the standard deviation for these six profiles. (B) C_α_ RMSD distributions (KDE) for the M2/M3 loop residues after superposition on the loop from the starting model (5VRF). (C) M2/M3 loop (gold) conformations for the apo simulation md1, protomer B. The image depicts 1000 frames from the trajectory taken at 1-ns intervals with individual frames superimposed on the C_α_ atoms of the TMD. The Zn-free 5VRF structure is shown in purple. (D) M2/M3 loop (gold) conformations for the holo simulation md1, protomer B (as in C); 5VRF starting conformation shown in cyan with Zn ions in magenta.

**Figure 6.**
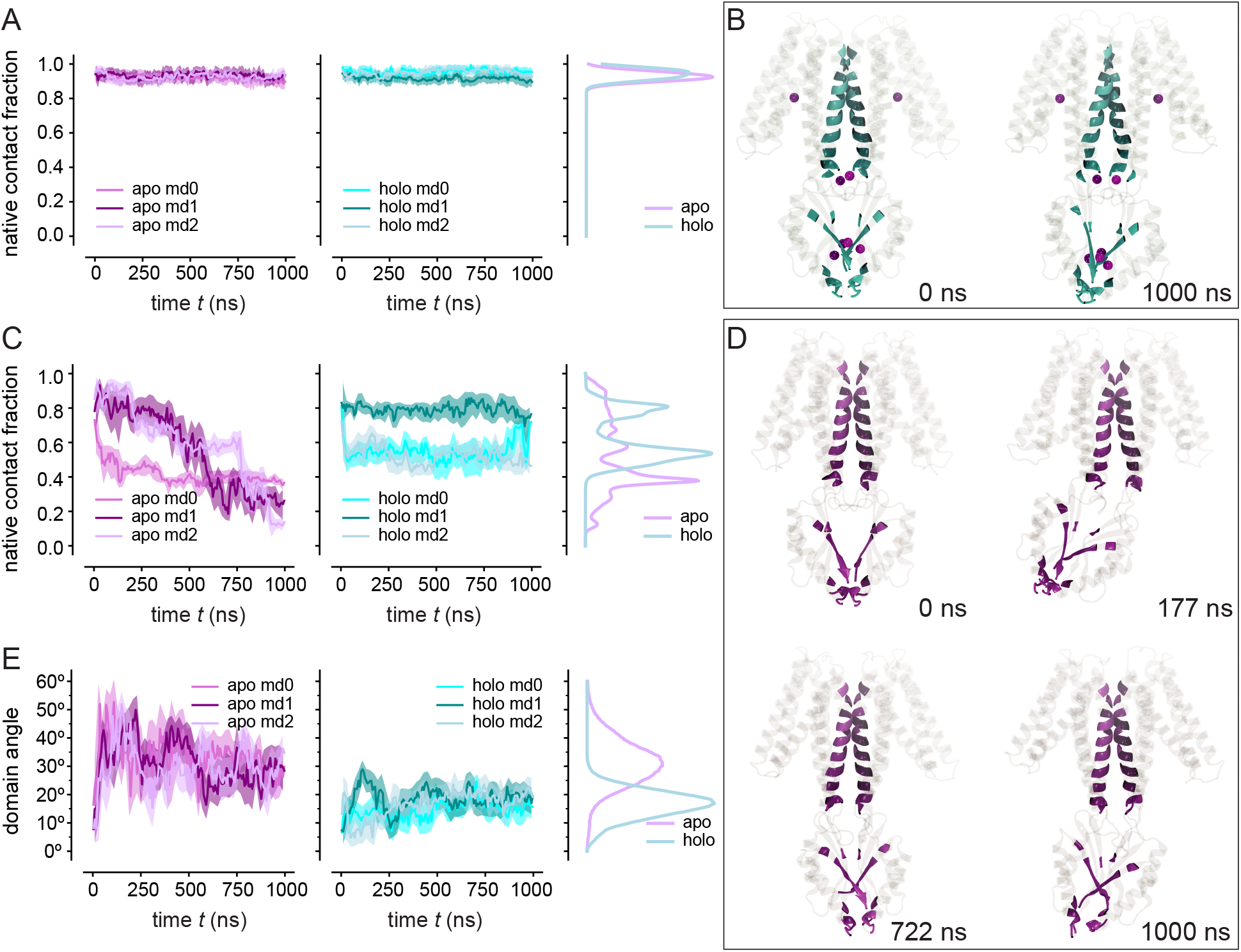
Stability of the dimer interface in MD simulations. (A) Dimeric interactions in the TMD quantified as the fraction of native contacts (from 5VRF) retained over the course of the simulations for apo (purple) and holo (cyan) states. Timeseries for all three repeats were averaged over 20 ns time intervals (solid line) with 95% confidence intervals shown as transparent bands. Distributions (KDEs) from all data are shown on the right. (B) Snapshots from a holo MD simulation (md1) at the indicated times showing the TMD native contact residues (K79, L83, L86, A87, A90, F91, M93, G94, S95, F97, L98, L99, Y102, E105, L207, L208) and CTD native contact residues (R237, G255, L257, S258, L259, N260, H263, E281, I283, I284, H285, Q286, D287, P288) in cyan. Magenta spheres represent Zn ions. The membrane, water, and other ions are omitted for clarity. (C) Dimeric interactions in the CTD as in panel A. (D) Snapshots from an apo simulation (md1) at the indicated times showing that dimeric interactions remain intact despite large interdomain movements. (E) Change in the angle between TMD and CTD from MD simulations. The angle corresponds to that required for optimal superimposition of the Cα atoms from the CTD to the reference structure (5VRF) at each time point in the MD trajectory. This angle was determined after aligning the TMD with the starting structure.

### Structural heterogeneity induced by metal ion chelation

Two additional data sets (denoted SP3 and SP4) were collected after chelation of metal ions with EDTA and/or TPEN in an attempt to improve the resolution of the apo state (Table I). However, classification of particles from each of these data sets resulted in two distinct structures corresponding to C2 symmetric and bent conformations that were seen in the SP1 and SP2 structures, respectively (Suppl. Figs. 9 & 10). In this way, SP3 and SP4 data sets generated four additional structures: SP3sym, SP3bent, SP4sym, and SP4bent (Table II). Due to the increased numbers of particles and perhaps to the use of the rigidified Fab2r construct, the resolutions of symmetric structures were improved (3.4 Å and 3.5 Å for SP3sym and SP4sym respectively). An atomic model was built and refined for the highest resolution structure (SP3sym) and comparison of this model with that from SP1 generated an RMSD of 1.2 Å, both for the YiiP monomer and dimer, indicating that the structures are essentially identical (Fig. 7a, Table III). Similarly, a model for the SP4sym map, built using Namdinator (Kidmose et al., 2019), produced RMSD’s of 1.4 Å and 1.3 Å relative to SP1 and SP3sym models, respectively. Despite overall similarity of these models, subtle differences were observed at the ion binding sites in the maps. In particular, SP1 and SP3sym maps had clear densities at all three Zn^2+^ binding sites, whereas SP4sym lacked density at site A in the membrane (Fig. 7b) and at site C in the CTD (Fig. 7c). In contrast, the M2/M3 loop was well ordered in all three maps with density visible at site B (Fig. 7d). This observation suggests that occupancy of site B may be the key to maintaining this symmetric conformation, whereas occupancy of sites A and C is secondary.

**Figure 7.**
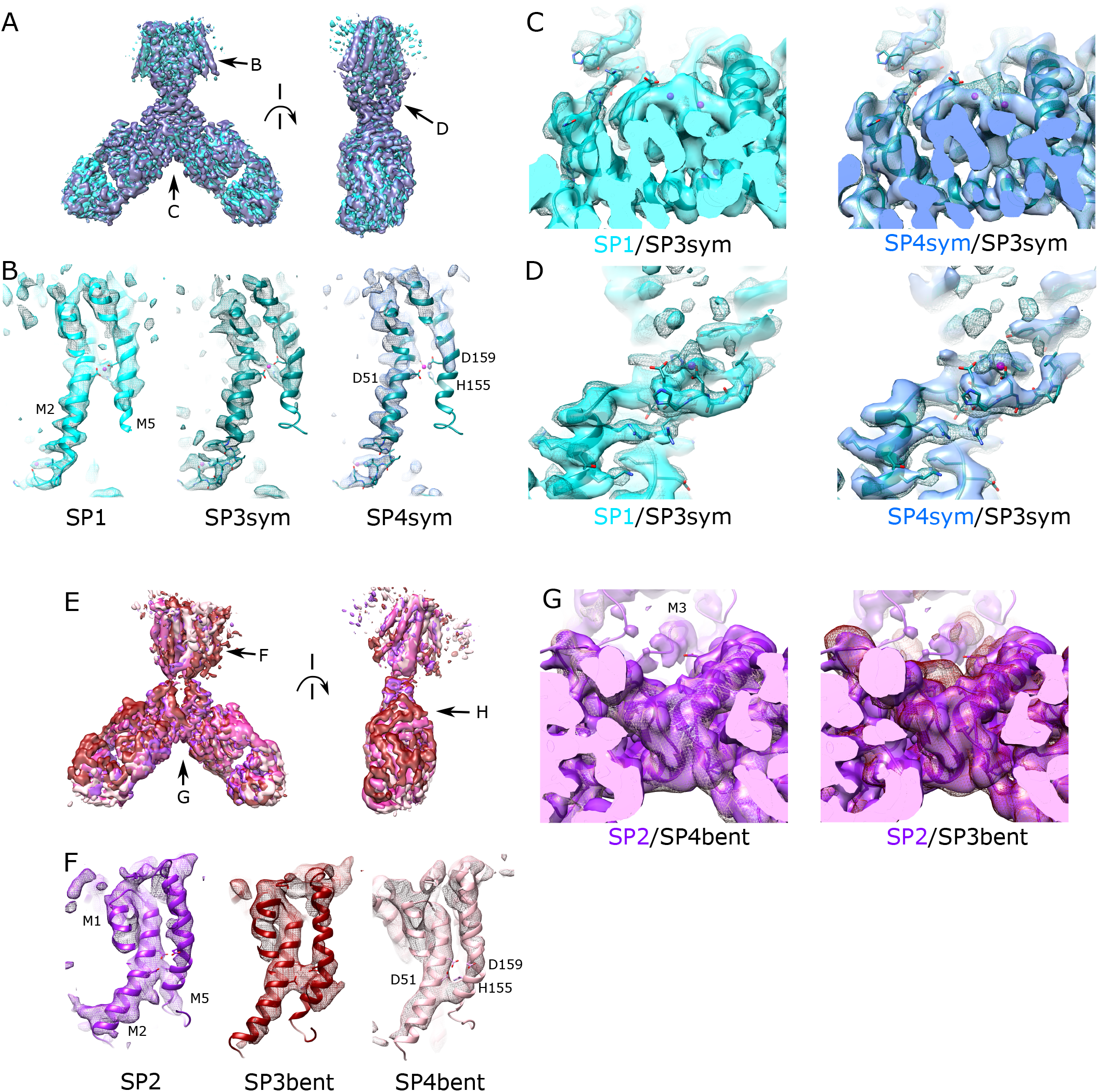
Comparison of apo and holo states derived from the various independent data sets. (A) Overlay of density maps from C2 symmetric conformations from SP1, SP3sym and SP4sym data sets; there is a 90° rotation between the two views. (B) Close-up view of Zn binding site A from the three data sets. Models for SP1 and SP3sym were independently refined and Namdinator was used to generate an MDFF model for SP4sym. Density is visible at the site in SP1 and SP3sym maps, but absent from the SP4sym map. (C) Comparison of site C in the CTD shows that extra density is present at site C in SP1 (solid cyan surface) and SP3sym (mesh surface) maps, consistent with the presence of metal ions. However, these densities are much weaker SP4sym map (solid blue surface). (D) Comparison of site B in the TM2/TM3 loop show that the loop is ordered and density associated with a metal ion is present in all three maps. (E) Overlay of density maps from the asymmetric conformations from SP2, SP3bent and SP4bent datasets; there is a 90° rotation between the two views. (F) Close-up view of Zn binding site A from the three data sets. The model for SP1 was manually built and refined, whereas MDFF models for SP3bent and SP4bent were generated by Namdinator. All density maps show a lack of density at the Zn site A. (G) Comparison of site C in the CTD shows a lack of density associated with metal ions in SP2 (solid purple surface) and SP4bent data sets (plum mesh). However, extra density is seen at this site in the map from SP3bent data set (dark red mesh).

A similar comparison of the asymmetric, bent structure from SP2 with those from SP3bent and SP4bent indicates that they all represent the same conformation (Fig. 7e). In contrast to the symmetric structures, the resolution of the bent structures were not improved relative to SP2 (4.2 and 4.7 Å for SP3bent and SP4bent, respectively). Given the lower resolution, we used Namdinator to build models for SP3bent and SP4bent maps based on the model from SP2. The resulting MDFF models had RMSD for C_α_ atoms of 1.5 Å and 2.0 Å relative to the SP2 model (Table III), respectively, indicating nearly identical conformations represented by these structures. Inspection of Zn^2+^ site A revealed a lack of density for the metal ion and a bent M2 helix in all of the maps (Fig. 7f). All of the maps also featured disordered M2/M3 loops indicating that Zn^2+^ site B was completely disrupted.

Careful inspection of CTD’s from the various maps suggests that Zn^2+^ binding at site C leads to a more compact dimer interface. At site C in the CTD, density for metal ions was lacking for SP2 and SP4bent maps; however, the map from SP3bent unexpectedly revealed at least partial density at this location (Fig. 7g). In the corresponding atomic models, distances between C_α_ atoms of Arg237 and Glu281 measured 11.1 Å for SP3bent compared to 16.0 Å for SP2 and SP4bent (Suppl. Fig. 11). An analogous difference was also evident in the overlay of CTD’s from SP1 and SP2 (12.2 Å vs. 16 Å, Fig. 4d). In fact, the compact spacing was seen in all of the maps showing density at site C (SP1, SP3sym, SP4sym, SP3bent) whereas the larger spacing was seen in maps lacking this density (SP2 and SP4bent, Suppl. Fig. 11). Although relatively subtle, this difference suggests that Zn^2+^ binding to site C produces a more compact dimer interface. This observation correlates with our MD simulations in which dimeric contacts in the CTD are gradually lost in the apo state (Fig. 6c) and the distance between Arg237 and Glu281 is considerably more variable (Suppl. Fig. 11c).

### Structural variability of YiiP

MD simulations and cryo-EM structures both indicate that the YiiP dimer has enhanced flexibility in the apo state. Specifically, the MD simulations starting from the symmetric holo state with Zn^2+^ removed displayed enhanced inter-domain movements and disordering of the M2/M3 loop. The cryo-EM structures of the apo state display lower resolution and disruption of the two-fold symmetry seen in previous structures of the holo state. Although our simulations of the apo state do not sample the fully bent conformation represented by the SP2 structure, the heterogeneity present in the SP3 and SP4 data sets provides an opportunity to visualize this transition. We used the 3D variability job in cryoSPARC (Punjani and Fleet, 2020) to visualize this flexibility and to map the transition between the two states. This non-linear analysis deduces principal components of the variability within a given particle population and produces a series of maps that represent the landscape of conformations represented by each component. We generated animations for the first three components - representing the highest variance - for both SP3 and SP4 data sets (Movie Suppl. 1-6). Although each animation shows unique aspects of heterogeneity, such as the pivoting of the distal Fab domain and general disordering of the TMD, the movements seen in component 1 from SP4 appear to capture the transition between the holo and apo states. Over the course of the corresponding animation (Movie Suppl. 5, Suppl. Fig. 12), a bend develops between the CTD and the TM domain which is accompanied by disordering of the M2/M3 loop. Also of interest is component 0 from this SP4 data set (Movie Suppl. 4) with movements of the CTD that reflect flexibility of the CTD dimer interface discussed above. Component 2 from the SP3 data set (Movie Suppl. 3) show the membrane domain swinging from side to side as the TM2/TM3 loops transition between order and disorder. This behavior is consistent with the idea that movements of the TMD relative to the CTD are coupled to Zn binding at site B. In order to evaluate these structural changes more objectively and compare them with the holo and apo state conformations, we used Namdinator to generate a series of models based on the maps from 3D variability analyses. This approach to model building has limited accuracy due to the down-sampling of maps for 3D variability analysis (5.5 Å), but the trajectory based on component 1 from SP4 clearly illustrated the molecular transition from the symmetrical, holo state to the asymmetrical apo state structures (Suppl. Fig. 12, Movie Suppl. 7). These models, which were generated objectively from SP3 and SP4 data sets, serve as an objective validation of the conformational transition represented by SP1 and SP2 data sets, featuring bending of M2 and M5 as well as disordering of the M2/M3 loop.

### Zn^2+^ binding controls a hydrophobic gate

Comparison of structures representing apo and holo states as well as the 3D variability analysis indicate that a hydrophobic gate on the cytoplasmic side of M5 and M6 appears to be controlled by Zn binding. Specifically, Leu154 on M5 and Leu199 on M6 are homologous to residues previously identified by X-ray-mediated hydroxyl radical labelling of YiiP from *E. coli* and proposed to control access to the transport sites (Gupta et al., 2014). In our structures, these residues are brought together during the transition from holo to apo state (Fig. 8). Although full closure occurs only on one protomer in the 3D variability analysis of SP4 (Suppl. Fig. 12, Movie Suppl. 7), the homogeneous data sets of the apo state (SP2, SP3bent and SP4bent) show closure of this gate on both protomers, with distances of 9.0 - 9.3 Å between C_α_ atoms from the two Leu residues (side chain atoms are considerably closer), compared to 12.8 Å in the SP3sym structure of the holo state. This distance is plotted for the Namdinator structures generated for the 3D variability analysis, together with the distance between Ala43 and Ala185, which appear to form a gate on the periplasmic side of the membrane (Suppl. Fig. 12d). Although these plots support the closure of the hydrophobic gate on the cytoplasmic side of the membrane, there is no change in the periplasmic gate, indicating that the molecule remains in the inward-facing conformation.

**Figure 8.**
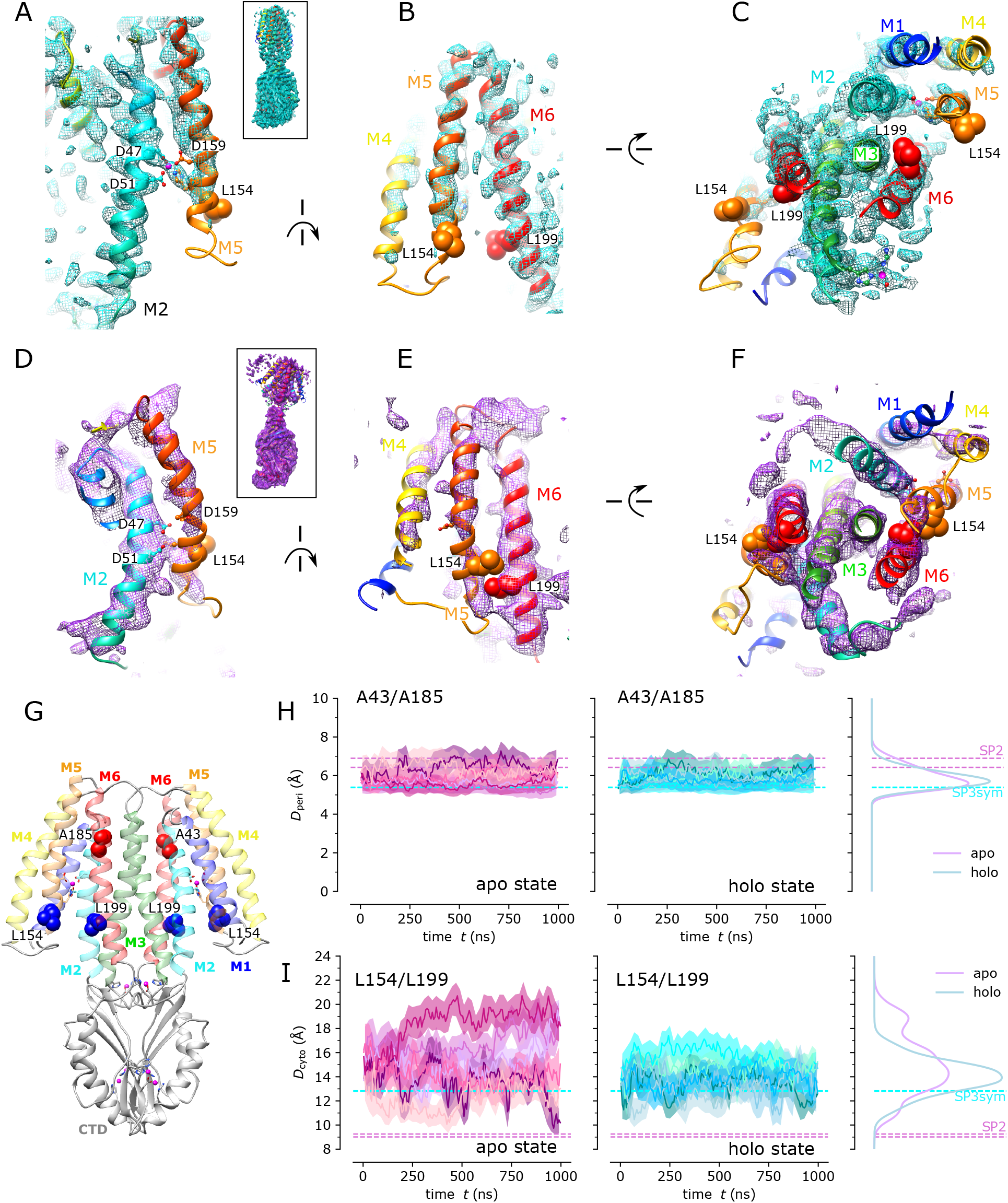
Zn binding controls a hydrophobic gate on the cytoplasmic side of the TMD. (A-C). Three orthogonal views of the cryo-EM structure of the symmetric holo state (SP3sym) showing clear separation between Leu154 and Leu199. Helices in the TMD have rainbow colors ranging from blue for M1 to red for M6. (D-F) Analogous views of the cryo-EM structure from the asymmetric apo state (SP2) showing how bending of M5 brings Leu154 and Leu199 close together. (G) Overview of the YiiP dimer showing the locations of residues used as collective variables: Leu154 and Leu199 are blue whereas Ala43 and Ala185 on the periplasmic side of the membrane are red. (H) Distances between C_α_ atoms of Ala43 and Ala185 (*D*_peri_) for each protomer during the three simulations in the apo (purple) and holo (cyan) states. Distributions calculated as KDEs are shown on the right. The corresponding values of *D*_peri_ from the SP3sym and SP2 structures are shown as dashed horizontal lines; SP2 is asymmetric and hence two values are shown. (I) Distances between C_α_ atoms of Leu154 and Leu199 (*D*_cyto_) for simulations of the apo and holo states with distributions shown on the right. *D*_cyto_ from the SP3sym and SP2 structures are shown as horizontal dashed lines.

In order to look for comparable movements in the MD simulations, we followed collective variables representing both cytoplasmic and periplasmic gates. Like the cryo-EM structures, the periplasmic gate was invariant in simulations of both apo and holo states (Fig. 8h). The cytoplasmic gate was considerably more dynamic, especially in the apo state. Although a stable closure did not occur, the distance between L154 and L199 dipped below 10 Å at several points during one of the simulations, ending at a value close to 9 Å (dark purple trace in Fig. 8i). This movement was only observed in one molecule, indicating that cooperativity between the monomers was weak. No change was observed in the distance between Ala43 and Ala185, consistent with the analysis of cryo-EM structures. Given the behavior of the periplasmic gate, the MD simulations offered no evidence of a conformational change to the outward-facing state.

## Discussion

In this study, we have examined the effect of Zn binding on the conformation and dynamics of YiiP from *S. oneidensis*. Like previous work with tubular crystals, we found metal ions bound to soYiiP despite their absence in buffers used for cell growth and protein purification. Metal ions remained bound even after transient treatment with chelator prior to 2D crystallization. These results indicate that soYiiP has a very high innate binding affinity at all three Zn sites and that ion binding is a prerequisite to formation of tubular crystals. To facilitate single-particle cryo-EM studies, an Fab was used to generate a complex and structures of the YiiP/Fab complex were resolved before and after treatment with the metal ion chelator EDTA. Although these data sets contained heterogeneous populations of molecules, we were able to generate structures of YiiP in holo, apo and partially bound states. Comparison of these structures revealed large scale conformational changes associated with the removal of metal ions. These changes included an ~25° bend of the TMD relative to the CTD, which broke the two-fold symmetry seen in previous structures (Coudray et al., 2013; Lopez-Redondo et al., 2018; Lu et al., 2009; Lu and Fu, 2007), bending of two key membrane helices and disordering of the M2/M3 loop, which harbors one of the Zn binding sites. Despite these dramatic changes, dimer interfaces in the TMD and CTD both remained intact and the molecule retained the inward-facing conformation in the absence of ions. Comparisons of refined structures, as well as principal components derived from 3D variability analysis, suggest that Zn binding at site B within the M2/M3 loop serves as a sensor that triggers this conformational change. Within the TMD, bending of M2 and M5 bring two hydrophobic residues together in what has previously been described as a hydrophobic gate, though access to the transport sites from the cytoplasm was not completely blocked and there was no evidence of a completed transition to the outward-facing conformation.

The hydrophobic gate was initially identified using hydroxyl radical footprinting to probe conformational changes of ecYiiP (Gupta et al., 2014). For this approach, EDTA-treated ecYiiP was exposed to an intense X-ray source to induce non-specific labelling of exposed residues, which were then identified by mass spectrometry. Addition of Zn to the buffer reduced labeling of residues that coordinate Zn at site A, as expected. In addition, the authors observed a significant decrease in labeling of Leu152 and Met197 which reside on the cytoplasmic end of M5 and M6, respectively. Based on this result, the authors postulated that Zn binding triggered the closure of this gate to produce an occluded state prior to the transport step. This report did not address the status of other Zn sites (B and C) nor did it address the starting conformation of ecYiiP prior to the addition of Zn. Although the current study confirms an interaction between these two hydrophobic residues (L154 and L199 in soYiiP), it suggests that the conformational dynamics are more complex. In particular, whereas hydroxy radical footprinting indicated that Zn closes the hydrophobic gate in ecYiiP, our cryo-EM study showed that the gate remained open in the holo state and that closure was elicited by chelation and removal of Zn. This discrepancy could reflect differing conformational equilibria of the two homologs, in which ecYiiP preferentially adopts the outward-facing state seen in the X-ray structures (Lu et al., 2009; Lu and Fu, 2007) whereas soYiiP prefers the inward-facing state seen in cryo-EM structures, both from membrane embedded tubular crystals and from detergent solubilized YiiP/Fab complexes. This discrepancy could reflect differing effects of Zn on these two states. Alternatively, the unwinding of TM helices and the splaying of TMD’s seen in the X-ray structures (Lopez-Redondo et al., 2018) may reflect an instability in the TM domain of ecYiiP such that the environment - lipid bilayer vs. detergent micelle - may play an important role in determining the conformational effects of Zn binding to the protein.

Comparison of the various structures produced from the heterogeneous data sets SP3 and SP4 suggest that Zn site B is the trigger for the structural difference between apo and holo states. More specifically, the symmetric structures from the SP1, SP3sym data sets show density at site A, but the SP4sym data set does not (Fig. 7b). Asymmetric structures from SP2, SP4bent data sets show a lack of density at site C, but the SP3bent data set appears to have retained ions at this site (Fig. 7g). Furthermore, the SP3bent structure retains the compact CTD dimer interface that characterizes the holo state (Suppl. Fig. 11). Thus, the status of site B is the only consistent difference between symmetric and bent conformations. Specifically, the M2/M3 loop is ordered with density visible at the binding sites in all the symmetric structures, whereas the loop is consistently disordered in the bent structures. Although this disorder makes it impossible to evaluate the presence of an ion, it seems unlikely that the coordinating residues (D70, H73, H77) would be able to form a site in the disordered state. The MD simulations are also consistent with these experimental observations, showing that the absence of Zn abolishes the binding site (Suppl. Fig. 4b) and greatly enhances the dynamics of this loop as well as movements of the CTD relative to the TMD.

In earlier reports, a role for site B in transport by YiiP has been largely overlooked, due to its lack of conservation among the family of CDF transporters (Kolaj-Robin et al., 2015). However, recent structural work on the mammalian homolog Znt8 revealed an unexpected Zn binding site that involves His137 in the M2/M3 loop (Xue et al., 2020). This site was readily titratable and the authors suggested that it might play a role in funneling ions into the cavity leading to the transport site. More generally, His-rich cytoplasmic lops are a relatively common feature of CDF transporters, though they often appear in other locations in the primary structure (Cotrim et al., 2019). In particular, the M4/M5 loop is highly enriched in His and Asp residues in a number of transporters from plants and yeast (Kawachi et al., 2008; Podar et al., 2012). The loop from AtMTP1 was shown to bind Zn and physiological studies of truncation mutants suggested that it functioned as a Zn sensor in a cellular context (Tanaka et al., 2015).

On the other hand, there is little evidence supporting a functional role for the binuclear Zn site C in the CTD. In our study, cryo-EM structures as well as MD simulations indicate that removal of Zn from this site enhances the flexibility of the dimer interface between the CTD’s. The resulting movements are, however, modest and the overall architecture of this domain remains unchanged. In the CTD of Znt8, one of the Zn ions is coordinated by conserved residues His234, His250, and Asp287 (site C2, Suppl. Fig. 4d). Although residues coordinating the second ion bound by the CTD of YiiP (site C1, Asp287, His263, and His285) are not conserved, the CTD of Znt8 unexpectedly bound a second ion via a unique CHC motif in a non-conserved N-terminal extension. Like our studies of YiiP, ions were observed bound to the CTD of Znt8 even when Zn^2+^ was not present in purification buffers, indicating high affinity and poorly exchangeable sites. Constitutive binding is also consistent with results from ITC binding studies with ecYiiP (Chao and Fu, 2004). In contrast, a structural study of isolated CTD’s from CzrB showed that ions were readily exchangeable and that binding was associated with a dramatic conformational change (Cherezov et al., 2008). However, the CTD construct used for this study lacked a conserved salt bridge between protomers at the membrane surface as well as the entire TMD, which likely serve as powerful constraints on CTD conformation and thus on the innate affinity of site C.

Although our refined structures of apo and holo states show both protomers in an equivalent conformation, the 3D variability analysis and MD simulations both suggest that each protomer can function independently. In particular, component 1 from the SP4 data set shows conformational changes in M2, M5 and the M2/M3 loop in only one of the protomers. Similarly, collective variables from MD simulations show that one protomer can close the hydrophobic gate at the cytoplasmic side of the TMD independently of its partner. This non-cooperative behavior is consistent with the structural results from Znt8, which revealed a dimer with mixed conformations for the WT protein prepared in the presence of Zn^2+^ (Xue et al., 2020). Although this structure was at lower resolution, it clearly showed movements of M1, M2, M4 and M5 that convert the outward-facing state to the inward-facing state in one of the protomers. Interestingly, the dimer interface mediated by M3 and M6 is unaffected by these movements, supporting results from cysteine crosslinking of residues along M3 that also imply that this interface remains static during transport (Lopez-Redondo et al., 2018). In Znt8, residues governing the hydrophobic gate are V219 in M5 and I266 in M6, which are in close apposition in the outward-facing state, but not in the inward-facing state. Using this structure as a template for the endpoints of the transport cycle, one could imagine that the apo conformation that we have observed is a first step in the process, which involves only M2 and M5. A second step, involving tilting of M1 and M4, would be required to complete the transition. Clearly the individual transporters have different conformational energy landscapes that are likely to reflect the different physiological environments in which they function. In future studies, effects of the environment used to preserve the membrane domain (e.g., lipid bilayer vs. detergent micelle) should be explored as they may hold the key to a better understanding of the conformational changes that govern the transport cycle.

## Acknowledgments

Funding for this work was provided by NIH grant 1R01GM125081 to DLS.

Electron microscopy was performed at the Cryo-EM Core Facility at NYU Langone Health, with the assistance of William Rice and Bing Wang. In addition, electron microscopy was performed at the Pacific Northwest Center for Cryo-EM at Oregon Health Sciences University, which was supported by NIH grant U24GM129547 and accessed through EMSL (grid.436923.9), a DOE Office of Science User Facility sponsored by the Office of Biological and Environmental Research.

MD simulations were performed using PSC Bridges at the Pittsburgh Supercomputing Center (allocation TG-MCB130177), a resource of the Extreme Science and Engineering Discovery Environment (XSEDE), which is supported by National Science Foundation grant number ACI-1548562. The authors also acknowledge Research Computing at Arizona State University for providing HPC and storage resources that have contributed to the research results reported within this paper.

## Methods

### Protein expression and Purification

Wild-type YiiP was expressed in *E. coli* BL21-AI cells (Life Technologies) from a modified pET vector with an N-terminal decahistidine tag. Cells were grown in LB media supplemented with 30 μg/ml kanamycin at 37°C until they reached an A_600_ of 0.8, at which point they were rapidly cooled to 18°C. Expression was induced by addition of 0.1 mM IPTG followed by overnight incubation at 18°C. Cells were harvested by centrifugation at 4,000xg for 1 h, resuspended in lysis buffer (20 mM Hepes pH 7.5, 100 mM NaCl, 10% glycerol, 250 μM TCEP) and then lysed with a high-pressure homogenizer (Emulsiflex-C3, Avestin). The membrane fraction was collected by centrifugation at 100,000xg and then solubilized by addition of 0.375 g n-dodecyl-β-D-maltoside (DDM, Anatrace) per gram of the membrane pellet for 2 h at 4°C in lysis buffer. Insoluble material was removed by centrifugation at 100,000xg for 40 min. The supernatant was loaded onto a Ni-NTA affinity column pre-equilibrated in buffer A (20 mM Hepes pH 7.5, 100 mM NaCl, 10% glycerol, 0.05% DDM). The column was washed by addition of Buffer A supplemented with 20 mM imidazole and protein was then eluted using a gradient of imidazole ranging from 20 - 500 mM. Peak fractions from this elution were combined and dialyzed overnight at 4°C against buffer A. During this dialysis, the decahistidine tag was cleaved by inclusion of TEV protease (1:10 weight ratio of TEV:YiiP). TEV was removed by loading the dialysate onto a Ni-NTA column and collecting the flow-through fractions. After concentration, a final purification was done with a Superdex 200 SEC column (GE Healthcare) equilibrated with SEC buffer (20 mM Hepes pH 7.5, 150 mM NaCl, 0.2% n-decyl-β-D-maltoside [DM], 1 mM TCEP).

### Transport assay

In order to measure transport, YiiP was reconstituted into proteoliposomes for a fluorometric assay described previously (Lopez-Redondo et al., 2018). For reconstitution, 25 μg of purified protein in reconstitution buffer (20 mM Hepes pH 6.5, 200mM K_2_SO_4_) was mixed with 1.5 mg Triton X-100 and 2.5 mg of *E. coli* polar lipids (EPL, Avanti Polar Lipids) in a volume of 250 μl. This protein/lipid/detergent mixture was incubated at room temperature for 30 min and then SM2 BioBeads (BioRad) were added in 3 steps: 7.5 mg BioBeads followed by 1.5 h incubation at room temperature, 15 mg BioBeads followed by overnight incubation at 4°C, and 15 mg BioBeads for 1hr at 4°C. The resulting proteoliposomes were collected and stored at −80°C.

Prior to the transport assay, proteoliposomes were loaded with 200 μM of FluoZin-1 (ThermoFisher Scientific) by 5 cycles of freeze-thaw using LN_2_. The proteoliposomes were then extruded 13 times through 0.4 μm polycarbonate membranes (Whatman Nucleopore, Millipore Sigma) pre-equilibrated with reconstitution buffer; excess FluoZin-1 dye was removed by passing the sample through a PD-10 desalting column (GE Healthcare) pre-equilibrated with reconstitution buffer.

For the transport assay, proteoliposomes were introduced into a fluorimeter (Fluoromax-4, Horiba Scientific) using a stopped-flow apparatus (STA-20 Rapid Kinetic Accessory, Hi-Tech Scientific) with a dead time of ~50 msec. This apparatus mixed the proteoliposome solution with an equal volume of reconstitution buffer supplemented with various concentrations of ZnCl_2_ (0.125mM – 64 mM). Fluorescence was excited at 490 nm and monitored at 525 nm. To normalize the fluorescence signal, the maximal fluorescence from each individual preparation was determined after adding 4% β-octyl-D-glucoside (OG) and 64 mM ZnCl_2_ to the proteoliposomes. In addition, a protein-free preparation of liposomes was used to determine background signal. From these data, the transport rate was quantified by plotting FP/FPmax − FL/FLmax vs. time, where FP and FL are the signals from proteoliposomes and protein-free liposomes, respectively, and FPmax and FLmax are the corresponding normalization signals in the presence of OG. The initial rates from each run were then fitted with the Hill equation to determine K_0.5_, n and V_max_ (Lopez-Redondo et al., 2018).

### Fab selection, expression and mutagenesis

Synthetic antibodies in the Fab format recognizing YiiP were selected from a phage-display library (Miller et al., 2012; Sauer et al., 2020). For this screen, purified YiiP retaining the His tag was reconstituted into biotinylated nanodiscs (Ritchie et al., 2009). For reconstitution, we used a construct of the membrane scaffolding protein (MSP) based on MSP1E3D1 with a Cys residue engineered into the C-terminus. This Cys residue was labeled with biotin by incubation of MSP with a 20-fold molar excess of maleimide-PEG11-biotin (ThermoFisher Scientific) overnight at 4°C. To reconstitute YiiP into nanodiscs, EPL was solubilized in DDM (1:2 weight ratio) and added to YiiP at a molar lipid-to-protein ratio of 277:1 in nanodisc buffer (20 mM Tris pH 7.4, 100 mM NaCl, 0.5 mM EDTA, 0.5 mM TCEP). After incubation on ice for 10 min, biotinylated MSP was added at an 8-fold molar excess to produce 500 μl of a solution containing 2.5 mM EPL, 72 μM MSP, 9 μM YiiP and 7.2 mM DDM. After incubation at 4°C for 1 h, BioBeads were added in three steps - 300 mg for 1 hr, 200 mg for 30 min and 100 mg overnight - while the solution was gently stirred at 4°C. Nanodiscs containing YiiP were then separated from empty nanodiscs by incubating the solution with 0.5 ml of Ni-NTA beads pre-equilibrated with nanodisc buffer for 30 min at 4°C. After packing these beads into a column and washing with 1 ml of nanodisc buffer, nanodiscs containing YiiP were eluted with 2 ml nanodisc buffer supplemented with 0.3 M imidazole. Peak elution fractions were pooled and concentrated with an Amicon concentrator (cutoff 50KDa). The concentrated sample was fractionated on Superdex 200 10/300 GL size exclusion column pre-equilibrated and eluted with nanodisc buffer.

For the phage display selection, biotinylated nanodiscs were immobilized to streptavidin-coated magnetic beads. Phage-display library sorting was performed as published previously (Dominik and Kossiakoff, 2015; Miller et al., 2012; Sauer et al., 2020). The process consisted of four rounds of binding, washing, and amplification of the bound phages in *E. coli*. In later rounds, enriched phage pools were reapplied to the bait either to improve the selection or to perform a negative selection against empty nanodiscs prepared in the absence of YiiP. Clones were analyzed using phage ELISA. The Fab genes of selected clones were subcloned into the Ptac-Fab-accept vector that was constructed from the RH2.2IPTG vector (a gift of Dr. Sachdev Sidhu) for large scale expression. After selecting a particular Fab molecule for cryo-EM analysis (see below), a mutation was made to the sequence of the hinge between the two domains of the Fab heavy chain in order to produce a more rigid molecules. Specifically, the sequence ^130^SSASTKG^136^, which is an invariate part of the Fab scaffold, was changed to ^130^FNQIKG^135^ (Bailey et al., 2018).

For expression of Fab, *E. coli* strain 55244 was transformed with the Fab expression vector. These cells were then cultured in 1 L TBG/Ap150 media ((12 g/L tryptone, 24 g/L yeast extract, 12.5 g/L K_2_HPO_4_, 2.3 g/L KH_2_PO_4_ and 0.8% glycerol) for ~24 h at 30°C. Although this plasmic carries a T4 promoter, Fab expression was achieved without addition of IPTG. Cells were harvested by centrifugation at 8,000xg for 1 h and the cell pellet was typically stored at −80°C. For purification, the cell pellet was thawed and resuspended in 20 mM Na_3_PO_4_ pH 7, with 1 mg/ml hen egg lysozyme, 1 mM PMSF and 1 μg/ml DNaseI/MgCl_2_. This cell suspension was incubated at room temperature for 1 h followed by 15 min on ice and then was passed through a high-pressure homogenizer (Emulsiflex-C3, Avestin). The lysate was centrifuged at 34,000xg for 1 h at 4°C and the supernatant passed through a 0.22 μm filter. This solution was loaded onto a 5 ml HiTrap Protein G HP column (GE Healthcare) equilibrated with 20 mM Na_3_PO_4_ pH 7 and eluted with 0.1 M glycine-HCl pH 2.7. Fractions of 2 ml were collected in tubes containing 200 μl of 2 M Tris-HCl pH 8. Pooled fractions were dialyzed against 1 L of 50 mM NaCH_3_CO_2_ pH 5 overnight at 4°C. Finally, this dialysate was loaded onto a Resource-S cation exchange column (GE Healthcare) equilibrated in 50 mM NaCH_3_CO_2_ pH 5 and eluted with a 0-50% gradient of elution buffer (50 mM NaCH_3_CO_2_ pH 5, 0.5 M NaCl). Fractions containing pure Fab were pooled and dialyzed against YiiP SEC buffer.

### Cryo-EM sample preparation and data analysis

A complex between YiiP and Fab was produced by incubating a mixture of the two proteins at a 1:1 molar ratio for 1 h at 20°C with total protein concentration of ~17 μM. For initial characterization of the complex, the sample was run on a Shodex KW-803 SEC column (Showa Denko America, Inc), which was equilibrated with 20 mM Hepes pH 7.5, 150 mM NaCl, 0.2% DM, 1 mM TPEN using an HPLC (Waters Corp, Milford MA) with a flow rate of 0.5 ml/min. The complex was then purified on a Superdex 200 SEC column equilibrated with SEC buffer using an FPLC (AKTA, GE Healthcare). Some of the samples were treated with either EDTA or TPEN in order to chelate metal ions (Table I). In those cases, the complex was incubated with 0.5 mM EDTA or with a combination of 0.5 mM EDTA and 0.5 mM TPEN for 16 h at 4°C and in most cases those chelators were also added to the SEC buffer during final purification. The peak elution fractions were pooled and concentrated to 3-4 mg/ml and used immediately for preparation of cryo-EM samples. For this process, 3–4 μl of solution were added to a glow-discharged grids (C-Flat 1.2/1.3-4Cu-50, Protochips, Inc) which were blotted under 100% humidity at 4°C and plunge frozen in liquid ethane using a Vitrobot (FEI Corp).

For tubular crystals, we followed procedures described in previous publications (Coudray et al., 2013; Lopez-Redondo et al., 2018). In brief, purified protein containing 0.9 mg/ml YiiP and 0.2% DM was mixed with a solution of 2 mg/ml dioleoylphosphatidylglycerol (DOPG) solubilized with 4 mg/ml DDM to achieve a lipid-to-protein weight ratio of 0.5. After incubation for 1 h, this solution was dialyzed for 14 d at 4°C against a buffer composed of 20 mM TES pH 7, 100 mM NaCl, 5 mM MgCl_2_ and 5 mM NaN_3_. The presence of tubular crystals was confirmed by viewing negatively stained samples. For cryo-EM analysis, samples were diluted 30-fold and applied to grids covered with home-made lacey carbon films. These grids were then blotted with filter paper applied to the back side of the grid and plunge frozen in liquid ethane using a Leica EMGP (Leica Microsystems).

Samples of tubular crystals were imaged with a Talos Arctica 200 kV electron microscope (FEI Corp) with a K2 Summit detector (Gatan Inc) and samples of the YiiP/Fab complex were imaged with a Titan Krios G3i, 300kV electron microscope (FEI Corp) equipped a Bioquantum energy filter with K2 or K3 direct electron detector (Gatan Inc). Pixel size was ~1 Å/pix with a total dose of 50-80 electrons/Å^2^. After selecting micrographs free from excessive contamination, crystalline ice or imaging artifacts, images of the YiiP/Fab complex were evaluated using cryoSPARC v2.15 (Punjani et al., 2017). Templates for particle picking were initially produced by manually picking ~1000 particles; 2D class averages were then used to identify particles in all micrographs. The initial set of particles were subjected to 2D classification followed by successive rounds of *ab initio* reconstruction with C1 symmetry and 2 classes to select a homogenous population. These particles were then used for heterogeneous refinement against 2 or 3 reference structures derived from the ab initio jobs. A final selection of particles was then used for non-uniform refinement with either C1 or C2 symmetry. Post processing steps included calculation of local resolution and evaluation of 3D variability (Punjani and Fleet, 2020). Images of the tubular crystals were analyzed using Relion according to protocols previously described (Lopez-Redondo et al., 2018).

For model building, we started with the deposited PDB files for YiiP (PDBID 5VRF) as well as a related Fab molecule (PDBID 4JQI). For Fab, we used MODELLER (Webb and Sali, 2014) to produce a homology model for Fab2. After rigid-body docking of these starting models to the map from SP1, we used Namdinator (Kidmose et al., 2019) to apply MDFF (Trabuco et al., 2008) followed by standard real-space refinement with PHENIX (Adams et al., 2010). For SP1, SP2, SP3sym data sets, we also used Coot (Emsley et al., 2010) to manually adjust the models to resolve errors or delete disordered loops followed by further rounds of real-space refinement with PHENIX. This process was repeated until acceptable refinement metrics were obtained. The model for the SP1 data set was used as the starting point for model building into the higher resolution map from the SP3sym dataset. For the SP2 dataset, the membrane domain was manually adjusted to match the map. Lower resolution in this region made it difficult to define the structures for some loops and membrane helices. For these regions, we used Namdinator models that were generated from the 3D variability analyses. In particular, after running the 3D variability job in cryoSPARC to generate 3 principle components, we used the “simple mode” for a 3D variability display job to produce 20 intermediate maps for each component. We then docked the model from SP3sym to the 10th map and used Namdinator for automated fitting. The resulting model was then used for Namdinator fitting to the 9th map and the resulting model was used for Namdinator fitting to the 8th map, etc. This iterative process was repeated to produce models for the 11th through 20th maps. Thus, this procedure objectively generated a model for the bent conformation, which contained bends in M2 and M5 as well as a plausible structure for the disordered M2/M3 loop. This Namdinator model was then docked to the SP2 map and used for several rounds of Coot modeling and PHENIX real-space refinement.

### Model for MD simulations

The cryo-EM structure of the zinc transporter YiiP from helical crystals (PDBID 5VRF) was used as the initial structure for all-atom MD simulations. All ionizable residues were set to their default protonation states. We determined the protonation states of histidines in the binding sites based on their orientation to Zn ions in the cryo-EM structure. H73 and H155 were modeled with the neutral “HSE” tautomer (proton on the N_ε_) while all other histidines were modeled with the neutral HSD tautomer (proton on N_δ_). As the cryo-EM structure contained zinc ions in its binding sites, it was directly used as the Zn^2+^-bound (holo) model. The starting model for the apo state was generated by simply removing the Zn ions from 5VRF.

### MD simulations

All-atom YiiP membrane-protein systems were built up in a 4:1 palmitoyloleoylphosphatidylethanolamine (POPE):palmitoyloleoylphosphatidylglycerol (POPG) bilayer, which approximates the composition of an *E. coli* bacterial plasma membrane (Raetz, 1986), with a free NaCl concentration of 100 mM using CHARMM-GUI v1.7 (Jo et al., 2008; Jo et al., 2009; Lee et al., 2016) with the CHARMM36 force field with the CMAP correction for proteins (MacKerell et al., 1998; Mackerell et al., 2004) and lipids (Klauda et al., 2010) together with the CHARMM TIP3P water model. The size of apo systems was 117,394 atoms in hexagonal simulation cells (101 Å × 101 Å × 139 Å), and that of holo systems was 115,068 atoms in hexagonal simulation cells (101 Å × 101 Å × 135 Å). Three repeats of 1-μs simulations were run for apo and holo state, starting from the same initial system conformation but with different initial velocities.

Simulations were performed using GROMACS 2019.6 (Abraham et al., 2015) on GPUs. Before the production equilibrium MD simulations, the systems underwent energy minimization and a 3.75-ns 6-stage equilibration procedure with position restraints on protein and lipids, following the CHARMM-GUI protocol (Jo et al., 2008). All simulations were carried out under periodic boundary conditions at constant temperature (T = 303.15 K) and pressure (P = 1 bar). A velocity rescaling thermostat (Bussi et al., 2007) was used with a time constant of 1 ps, and protein, lipids and solvent were defined as three separate temperature-coupling groups. A Parrinello-Rahman barostat (Parrinello and Rahman, 1981) with time constant 5 ps and compressibility 4.6 × 10^−5^ /bar was used for semi-isotropic pressure coupling. The Verlet neighbor list was updated every 20 steps with a cutoff of 1.2 nm and a buffer tolerance of 0.005 kJ/mol/ps. Coulomb interactions under periodic boundary conditions were evaluated with the smooth particle mesh Ewald (SPME) method (Essman et al., 1995) under tinfoil boundary conditions with a real-space cutoff of 1.2 nm and interactions beyond the cutoff were calculated in reciprocal space with a fast Fourier transform on a grid with 0.12-nm spacing and fourth-order spline interpolation. The Lennard–Jones forces were switched smoothly to zero between 1.0 and 1.2 nm and the potential was shifted over the whole range and reduced to zero at the cutoff. Bonds to hydrogen atoms were constrained with the P-LINCS algorithm (Hess, 2008) (with an expansion order of four and two LINCS iterations) or SETTLE (Miyamoto and Kollman, 1992) (for water molecules). The classical equations of motions were integrated with the leapfrog algorithm with a time step of 2 fs.

### Analysis of molecular dynamics simulations

Simulation trajectories were analyzed with Python scripts based on MDAnalysis (Gowers et al., 2016). Probability densities of timeseries data were calculated as kernel density estimates (KDE) using *sklearn.neighbors.KernelDensity* in the scikit-learn package (Pedregosa et al., 2011) with a bandwidth of 0.2. Root mean square deviations (RMSD) were calculated for C_α_ atoms of the whole protein, TMD and CTD after optimally superimposing the atoms on the same C_α_ atoms of the cryo-EM structure with the qcprot algorithm (Liu et al., 2010) as implemented in MDAnalysis. Root mean square fluctuations (RMSF) of all C_α_ atoms were calculated by superimposing the protein on either all, only TMD, or only CTD C_α_ atoms. In order to quantify the flexibility of the M2/M3 loop in apo and holo states, we also calculated the C_α_ RMSD’s of the loop when the whole protein was superimposed on itself or when just the loop was fitted on itself. To quantify the relative motion between the two domains, we performed a CTD-TMD rotation angle analysis. The whole protein was first superimposed on the TMD domain of a reference structure - here we used the experimental holo structure (PDBID 5VRF) as reference - and the rotation angle that minimized the RMSD of the CTD was calculated. To study the influence of Zn ions on the rigidity of the binding sites, RMSD’s of the side chains of the binding residues were calculated. The opening of the hydrophobic gates in the TMD was assessed using two collective variables: the distance between C_α_ atoms of Ala41 and Ala183 as the periplasmic gate and the distance between C_α_ atoms of Leu154 and Leu199 for the cytosolic gate. Native contacts analysis was used to describe the stability of dimeric interfaces in the TMD and CTD. The reference native contacts were defined as pairs of atoms in the TMD (primarily located in M3 and M6) or in the CTD between protomer A and protomer B, whose pair distance was shorter than 4.5 Å in the cryo-EM structure, 5VRF. A soft cut off (Best et al., 2013) with a softness of parameter of 5 Å^−1^ and a reference distance tolerance of 1.8 was used for the native contacts calculation, as implemented in MDAnalysis. Additionally, the RMSD timeseries of the native contact atoms were calculated. Additionally, the distance between C_α_ atoms of R237 and E281 was used as another collective variable to describe the CTD dimeric interface.

### Classical force field model for Zn(II) ions

Divalent ions are challenging to simulate with classical force fields, especially when ions need to be able to transition between solution and binding sites, as for the transport site A and potentially also for the B site in YiiP. Because only a simple non-bonded soft-sphere model for zinc was available in the CHARMM forcefield, we developed a 6-site non-bonded dummy model (Aqvist and Warshel, 1990; Duarte et al., 2014) for the Zn(II) ion in combination with the CHARMM TIP3P water model. Following the previous study (Duarte et al., 2014), the number of dummy sites (six) was selected based on the experimental coordination number of Zn ions in water (Marcus, 1988), and the model was optimized by reproducing the hydration free energy (HFE) of −1955 kJ/mol (Marcus, 1991) and an ion-oxygen distance (IOD) of 2.08 Å (Marcus, 1988) in simulations with the CHARMM TIP3P water model (see Suppl. Fig. 2). Each of the dummy sites (DM) was assigned a mass of 3u and a partial charge of δ=+0.35*e* while the central site (ZND) was assigned a mass of 47.39u and charge of –0.1*e* for a total charge of +2*e*, where *e* is the absolute value of the electron charge and u the atomic mass unit. Dummy atoms were constrained to be located at a fixed distance of 0.9 Å from the central atom; the overall arrangement of the dummy atoms at the corners of an octahedron was maintained by harmonic restraints between dummy atoms (force constant 334720 kJ/(mol nm^2^)) and angle restraints (force constant 1046 kJ/(mol rad^2^)). A genetic algorithm (Eiben et al., 1994) was used to optimize the Lennard-Jones parameters, which model van der Waals interactions, of both atom types. We first randomly generated eight sets of parameters. These sets were ranked based on the computed HFEs and the IODs. The HFEs were calculated via stratified all-atom alchemical free energy perturbation (FEP) MD simulations (Fan et al., 2020; Klimovich et al., 2015) and a finite size correction (Reif and Hünenberger, 2011) was applied to the simulated free energy values. In particular, a total correction of –150.1 kJ/mol, consisting of a type C_1_ correction of –149.8 kJ/mol for the use of a Ewald method to evaluate electrostatics, a type C_2_ correction of –1.3 kJ/mol to correct for the artifactual constraint of vanishing average potential in the Ewald method, and a type D (PBC/LS) correction of +1.0 kJ/mol to adjust for the difference in the dielectric permittivity between the water model and real water, was added to obtain final HFE that could be compared to the experimental value. IODs were calculated from 15-ns equilibrium MD simulations in the *NPT* ensemble at standard conditions as the position of the first peak of the radial distribution function (RDF) between Zn ions and the oxygen atoms of all water molecules. The best four sets of parameters were kept in the next generation. Four new sets were generated by taking a random weighted average of the parameters of two sets from the previous generation. Together with another four randomly generated parameter sets, a new generation with twelve candidates were again simulated and ranked by HFEs and IODs. After repeating these steps, we obtained a converged set of van der Waals parameters in the sixth generation that reproduced the experimental coordination number in water (6), ion-oxygen distance (model: 2.088 Å, experiment: 2.08 Å) and hydration free energy (model: –1956 kJ/mol, exp: –1955 kJ/mol) (see Suppl. Fig. 2a). The optimal Lennard-Jones parameters for the dummy sites (DM) were ε=4.541995 × 10^−4^ kJ/mol, σ=4.99926 × 10^−2^ nm and ε=21.2999497535 kJ/mol, σ= 1.30117 × 10^−1^ nm for the central ZND site. The final parameters were deposited in the Ligandbook repository (https://ligandbook.org/) (Domański et al., 2017) with package ID 2934 as input files for GROMACS.

As discussed in the Results section, our CHARMM Zn(II) non-bonded dummy model was validated with MD simulations using experimental crystal structures for β-1,3-1,4-endo-glucanase (PDBID 1U0A, resolution 1.64 Å), stromelysin-1 (2USN, 2.20 Å) and the δ’ subunit of *E. coli* clamp-loader complex (1A5T, 2.20 Å). Three independent repeats of 200-ns simulations were performed for each structure. The simulation systems were built with CHARMM-GUI v1.7 and the simulations were run with GROMACS 2019.6. The simulation settings were the same as for the YiiP simulations, except that we used isotropic pressure coupling and two separate temperature-coupling groups: protein and solvent. RMSD’s of backbone atoms, all binding site residues and ligand atoms were calculated to assess the stability of the Zn-bound structures. Radial distribution functions between the zinc ion and binding site ligand atoms were compared with the distances from crystal structures using RDF.InterRDFs in Parallel MD Analysis (PMDA) (Fan et al., 2019).

## References

Abraham, M.J., Murtola, T., Schulz, R., Páll, S., Smith, J.C., Hess, B., and Lindahl, E. (2015). GROMACS: High performance molecular simulations through multi-level parallelism from laptops to supercomputers. SoftwareX 1--2, 19–25, 10.1016/j.softx.2015.06.001.

Adams, P.D., Afonine, P.V., Bunkoczi, G., Chen, V.B., Davis, I.W., Echols, N., Headd, J.J., Hung, L.W., Kapral, G.J., Grosse-Kunstleve, R.W., et al. (2010). PHENIX: a comprehensive Python-based system for macromolecular structure solution. Acta Crystallogr D Biol Crystallogr 66, 213–221, 10.1107/S0907444909052925.

Andreini, C., Banci, L., Bertini, I., and Rosato, A. (2006). Counting the zinc-proteins encoded in the human genome. J Proteome Res 5, 196–201, 10.1021/pr050361j.

Aqvist, J., and Warshel, A. (1990). Free-Energy Relationships in Metalloenzyme-Catalyzed Reactions - Calculations of the Effects of Metal-Ion Substitutions in Staphylococcal Nuclease. Journal of the American Chemical Society 112, 2860–2868.

Bailey, L.J., Sheehy, K.M., Dominik, P.K., Liang, W.G., Rui, H., Clark, M., Jaskolowski, M., Kim, Y., Deneka, D., Tang, W.J., et al. (2018). Locking the Elbow: Improved Antibody Fab Fragments as Chaperones for Structure Determination. J Mol Biol 430, 337–347, 10.1016/j.jmb.2017.12.012.

Best, R.B., Hummer, G., and Eaton, W.A. (2013). Native contacts determine protein folding mechanisms in atomistic simulations. Proc Natl Acad Sci U S A 110, 17874–17879, 10.1073/pnas.1311599110.

Bussi, G., Donadio, D., and Parrinello, M. (2007). Canonical sampling through velocity rescaling. Journal of Chemical Physics 126, 10.1063/1.2408420.

Chao, Y., and Fu, D. (2004). Thermodynamic studies of the mechanism of metal binding to the Escherichia coli zinc transporter YiiP. J Biol Chem 279, 17173–17180, 10.1074/jbc.M400208200.

Cherezov, V., Hofer, N., Szebenyi, D.M., Kolaj, O., Wall, J.G., Gillilan, R., Srinivasan, V., Jaroniec, C.P., and Caffrey, M. (2008). Insights into the mode of action of a putative zinc transporter CzrB in Thermus thermophilus. Structure 16, 1378–1388, 10.1016/j.str.2008.05.014.

Cotrim, C.A., Jarrott, R.J., Martin, J.L., and Drew, D. (2019). A structural overview of the zinc transporters in the cation diffusion facilitator family. Acta Crystallogr D Struct Biol 75, 357–367, 10.1107/S2059798319003814.

Coudray, N., Valvo, S., Hu, M., Lasala, R., Kim, C., Vink, M., Zhou, M., Provasi, D., Filizola, M., Tao, J., et al. (2013). Inward-facing conformation of the zinc transporter YiiP revealed by cryoelectron microscopy. Proc Nat Acad Sci 110, 2140–2145, 10.1073/pnas.1215455110.

Cubillas, C., Vinuesa, P., Tabche, M.L., and Garcia-de los Santos, A. (2013). Phylogenomic analysis of Cation Diffusion Facilitator proteins uncovers Ni2+/Co2+ transporters. Metallomics 5, 1634–1643, 10.1039/c3mt00204g.

Domański, J., Beckstein, O., and Iorga, B.I. (2017). Ligandbook: an online repository for small and drug-like molecule force field parameters. Bioinformatics 33, 1747–1749, 10.1093/bioinformatics/btx037.

Dominik, P.K., and Kossiakoff, A.A. (2015). Phage display selections for affinity reagents to membrane proteins in nanodiscs. Methods Enzymol 557, 219–245, 10.1016/bs.mie.2014.12.032.

Duarte, F., Bauer, P., Barrozo, A., Amrein, B.A., Purg, M., Aqvist, J., and Kamerlin, S.C.L. (2014). Force Field Independent Metal Parameters Using a Nonbonded Dummy Model. Journal of Physical Chemistry B 118, 4351–4362, 10.1021/jp501737x.

Eiben, A.E., Raue, P.E., and Ruttkay, Z. (1994). Genetic algorithms with multi-parent recombination. Parallel Problem Solving from Nature - PPSN III - International Conference on Evolutionary Computation, Proceedings 866, 78–87.

Emsley, P., Lohkamp, B., Scott, W., and Cowtan, K. (2010). Features and development of Coot. Acta Crystallographica Section D - Biological Crystallography 66, 486–501.

Essman, U., Perela, L., Berkowitz, M.L., Darden, T., Lee, H., and Pedersen, L.G. (1995). A smooth particle mesh Ewald method. J Chem Phys 103, 8577–8592, 10.1063/1.470117.

Fan, S., Iorga, B.I., and Beckstein, O. (2020). Prediction of octanol-water partition coefficients for the SAMPL6-[Formula: see text] molecules using molecular dynamics simulations with OPLS-AA, AMBER and CHARMM force fields. J Comput Aided Mol Des 34, 543–560, 10.1007/s10822-019-00267-z.

Fan, S., Linke, M., Paraskevakos, I., Gowers, R.J., Gecht, M., and Beckstein, O. (2019). PMDA - Parallel Molecular Dynamics Analysis. In Proceedings of the 18th Python in Science Conference, C. Calloway, D. Lippa, D. Niederhut, and D. Shupe, eds. (Austin, TX: SciPy), pp. 134–142.

Gowers, R.J., Linke, M., Barnoud, J., Reddy, T.J.E., Melo, M.N., Seyler, S.L., Dotson, D.L., Domański, J., Buchoux, S., Kenney, I.M., et al. (2016). MDAnalysis: A Python package for the rapid analysis of molecular dynamics simulations. Paper presented at: Proceedings of the 15th Python in Science Conference (Austin, TX: SciPy).

Gupta, S., Chai, J., Cheng, J., D’Mello, R., Chance, M.R., and Fu, D. (2014). Visualizing the kinetic power stroke that drives proton-coupled zinc(II) transport. Nature 512, 101–104, 10.1038/nature13382.

Hess, B. (2008). P-LINCS: A Parallel Linear Constraint Solver for Molecular Simulation. J Chem Theory Comput 4, 116–122, 10.1021/ct700200b.

Jo, S., Kim, T., Iyer, V.G., and Im, W. (2008). CHARMM-GUI: a web-based graphical user interface for CHARMM. J Comput Chem 29, 1859–1865, 10.1002/jcc.20945.

Jo, S., Lim, J.B., Klauda, J.B., and Im, W. (2009). CHARMM-GUI Membrane Builder for Mixed Bilayers and its Application to Yeast Membranes. Biophys J 97, 50–58, 10.1016/j.bpj.2009.04.013.

Kambe, T. (2012). Molecular architecture and function of ZnT transporters. Current topics in membranes 69, 199–220, 10.1016/B978-0-12-394390-3.00008-2.

Kawachi, M., Kobae, Y., Mimura, T., and Maeshima, M. (2008). Deletion of a histidine-rich loop of AtMTP1, a vacuolar Zn(2+)/H(+) antiporter of Arabidopsis thaliana, stimulates the transport activity. J Biol Chem 283, 8374–8383, 10.1074/jbc.M707646200.

Kidmose, R.T., Juhl, J., Nissen, P., Boesen, T., Karlsen, J.L., and Pedersen, B.P. (2019). Namdinator - automatic molecular dynamics flexible fitting of structural models into cryo-EM and crystallography experimental maps. IUCrJ 6, 526–531, 10.1107/S2052252519007619.

Kim, C., Vink, M., Hu, M., Love, J., Stokes, D.L., and Ubarretxena-Belandia, I. (2010). An automated pipeline to screen membrane protein 2D crystallization. Journal of structural and functional genomics 11, 155–166, 10.1007/s10969-010-9088-5.

Kimura, T., and Kambe, T. (2016). The Functions of Metallothionein and ZIP and ZnT Transporters: An Overview and Perspective. Int J Mol Sci 17, 336, 10.3390/ijms17030336.

Klauda, J.B., Venable, R.M., Freites, J.A., O’Connor, J.W., Tobias, D.J., Mondragon-Ramirez, C., Vorobyov, I., MacKerell, J., Alexander D, and Pastor, R.W. (2010). Update of the CHARMM all-atom additive force field for lipids: validation on six lipid types. J Phys Chem B 114, 7830–7843, 10.1021/jp101759q.

Klimovich, P.V., Shirts, M.R., and Mobley, D.L. (2015). Guidelines for the analysis of free energy calculations. Journal of Computer-Aided Molecular Design 29, 397–411, 10.1007/s10822-015-9840-9.

Kolaj-Robin, O., Russell, D., Hayes, K.A., Pembroke, J.T., and Soulimane, T. (2015). Cation Diffusion Facilitator family: Structure and function. FEBS Lett 589, 1283–1295, 10.1016/j.febslet.2015.04.007.

Lee, J., Cheng, X., Swails, J.M., Yeom, M.S., Eastman, P.K., Lemkul, J.A., Wei, S., Buckner, J., Jeong, J.C., Qi, Y., et al. (2016). CHARMM-GUI Input Generator for NAMD, GROMACS, AMBER, OpenMM, and CHARMM/OpenMM Simulations Using the CHARMM36 Additive Force Field. J Chem Theory Comput 12, 405–413, 10.1021/acs.jctc.5b00935.

Li, P., and Merz, K.M., Jr. (2017). Metal Ion Modeling Using Classical Mechanics. Chem Rev 117, 1564–1686, 10.1021/acs.chemrev.6b00440.

Liang, X., Dempski, R.E., and Burdette, S.C. (2016). Zn(2+) at a cellular crossroads. Curr Opin Chem Biol 31, 120–125, 10.1016/j.cbpa.2016.02.008.

Liu, P., Agrafiotis, D.K., and Theobald, D.L. (2010). Fast determination of the optimal rotational matrix for macromolecular superpositions. J Comput Chem 31, 1561–1563, 10.1002/jcc.21439.

Lopez-Redondo, M.L., Coudray, N., Zhang, Z., Alexopoulos, J., and Stokes, D.L. (2018). Structural basis for the alternating access mechanism of the cation diffusion facilitator YiiP. Proc Natl Acad Sci U S A, 10.1073/pnas.1715051115.

Lovell, M.A., Smith, J.L., Xiong, S., and Markesbery, W.R. (2005). Alterations in zinc transporter protein-1 (ZnT-1) in the brain of subjects with mild cognitive impairment, early, and late-stage Alzheimer’s disease. Neurotox Res 7, 265–271, 10.1007/BF03033884.

Lu, M., Chai, J., and Fu, D. (2009). Structural basis for autoregulation of the zinc transporter YiiP. Nature structural & molecular biology 16, 1063–1067, 10.1038/nsmb.1662.

Lu, M., and Fu, D. (2007). Structure of the zinc transporter YiiP. Science 317, 1746–1748, 10.1126/science.1143748.

MacKerell, A.D., Bashford, D., Bellott, M., Dunbrack, R.L., Evanseck, J.D., Field, M.J., Fischer, S., Gao, J., Guo, H., Ha, S., et al. (1998). All-atom empirical potential for molecular modeling and dynamics studies of proteins. J Phys Chem B 102, 3586–3616, 10.1021/jp973084f.

Mackerell, A.D., Jr., Feig, M., and Brooks, C.L., 3rd (2004). Extending the treatment of backbone energetics in protein force fields: limitations of gas-phase quantum mechanics in reproducing protein conformational distributions in molecular dynamics simulations. J Comput Chem 25, 1400–1415, 10.1002/jcc.20065.

Marcus, Y. (1988). Ionic-Radii in Aqueous-Solutions. Chemical Reviews 88, 1475–1498, DOI 10.1021/cr00090a003.

Marcus, Y. (1991). Thermodynamics of Solvation of Ions .5. Gibbs Free-Energy of Hydration at 298.15-K. Journal of the Chemical Society-Faraday Transactions 87, 2995–2999, DOI 10.1039/ft9918702995.

Maret, W. (2013). Zinc biochemistry: from a single zinc enzyme to a key element of life. Adv Nutr 4, 82–91, 10.3945/an.112.003038.

Miller, K.R., Koide, A., Leung, B., Fitzsimmons, J., Yoder, B., Yuan, H., Jay, M., Sidhu, S.S., Koide, S., and Collins, E.J. (2012). T cell receptor-like recognition of tumor in vivo by synthetic antibody fragment. PLoS One 7, e43746, 10.1371/journal.pone.0043746.

Miyamoto, S., and Kollman, P.A. (1992). Settle - an Analytical Version of the Shake and Rattle Algorithm for Rigid Water Models. Journal of Computational Chemistry 13, 952–962, DOI 10.1002/jcc.540130805.

Montanini, B., Blaudez, D., Jeandroz, S., Sanders, D., and Chalot, M. (2007). Phylogenetic and functional analysis of the Cation Diffusion Facilitator (CDF) family: improved signature and prediction of substrate specificity. BMC genomics 8, 107, 10.1186/1471-2164-8-107.

Neupane, D.P., Kumar, S., and Yukl, E.T. (2019). Two ABC Transporters and a Periplasmic Metallochaperone Participate in Zinc Acquisition in Paracoccus denitrificans. Biochem 58, 126–136, 10.1021/acs.biochem.8b00854.

Parrinello, M., and Rahman, A. (1981). Polymorphic transitions in single crystals: A new molecular dynamics method. J~Appl~Phys 52, 7182–7190, 10.1063/1.328693.

Pedregosa, F., Varoquaux, G., Gramfort, A., Michel, V., Thirion, B., Grisel, O., Blondel, M., Prettenhofer, P., Weiss, R., Dubourg, V., et al. (2011). Scikit-learn: Machine Learning in Python. Journal of Machine Learning Research 12, 2825–2830.

Podar, D., Scherer, J., Noordally, Z., Herzyk, P., Nies, D., and Sanders, D. (2012). Metal selectivity determinants in a family of transition metal transporters. J Biol Chem 287, 3185–3196, 10.1074/jbc.M111.305649.

Prasad, A.S. (2013). Discovery of human zinc deficiency: its impact on human health and disease. Adv Nutr 4, 176–190, 10.3945/an.112.003210.

Punjani, A., and Fleet, D.J. (2020). 3D Variability Analysis: Resolving continuous flexibility and discrete heterogeneity from single particle cryo-EM. 2020.2004.2008.032466, 10.1101/2020.04.08.032466 %JbioRxiv.

Punjani, A., Rubinstein, J.L., Fleet, D.J., and Brubaker, M.A. (2017). cryoSPARC: algorithms for rapid unsupervised cryo-EM structure determination. Nat Methods 14, 290–296, 10.1038/nmeth.4169.

Raetz, C.R. (1986). Molecular genetics of membrane phospholipid synthesis. Annu Rev Genet 20, 253–295, 10.1146/annurev.ge.20.120186.001345.

Reif, M.M., and Hünenberger, P.H. (2011). Computation of methodology-independent single-ion solvation properties from molecular simulations. III. Correction terms for the solvation free energies, enthalpies, entropies, heat capacities, volumes, compressibilities, and expansivities of solvated ions. Journal of Chemical Physics 134, 10.1063/1.3567020.

Ritchie, T.K., Grinkova, Y.V., Bayburt, T.H., Denisov, I.G., Zolnerciks, J.K., Atkins, W.M., and Sligar, S.G. (2009). Chapter 11 - Reconstitution of membrane proteins in phospholipid bilayer nanodiscs. Methods Enzymol 464, 211–231, 10.1016/S0076-6879(09)64011-8.

Sala, D., Giachetti, A., and Rosato, A. (2019). An atomistic view of the YiiP structural changes upon zinc(II) binding. Biochim Biophys Acta Gen Subj 1863, 1560–1567, 10.1016/j.bbagen.2019.06.001.

Sauer, D.B., Trebesch, N., Marden, J.J., Cocco, N., Song, J., Koide, A., Koide, S., Tajkhorshid, E., and Wang, D.N. (2020). Structural basis for the reaction cycle of DASS dicarboxylate transporters. Elife 9, 10.7554/eLife.61350.

Sladek, R., Rocheleau, G., Rung, J., Dina, C., Shen, L., Serre, D., Boutin, P., Vincent, D., Belisle, A., Hadjadj, S., et al. (2007). A genome-wide association study identifies novel risk loci for type 2 diabetes. Nature 445, 881–885, 10.1038/nature05616.

Tanaka, N., Fujiwara, T., Tomioka, R., Kramer, U., Kawachi, M., and Maeshima, M. (2015). Characterization of the histidine-rich loop of Arabidopsis vacuolar membrane zinc transporter AtMTP1 as a sensor of zinc level in the cytosol. Plant Cell Physiol 56, 510–519, 10.1093/pcp/pcu194.

Trabuco, L.G., Villa, E., Mitra, K., Frank, J., and Schulten, K. (2008). Flexible fitting of atomic structures into electron microscopy maps using molecular dynamics. Structure 16, 673–683, 10.1016/j.str.2008.03.005.

Webb, B., and Sali, A. (2014). Protein structure modeling with MODELLER. Methods Mol Biol 1137, 1–15, 10.1007/978-1-4939-0366-5_1.

Xue, J., Xie, T., Zeng, W., Jiang, Y., and Bai, X.C. (2020). Cryo-EM structures of human ZnT8 in both outward- and inward-facing conformations. Elife 9, 10.7554/eLife.58823.

