## Supplemental Figures for "Zinc Dependent Conformational Changes in the Cation Diffusion Facilitator YiiP from S. oneidensis"

### Suppl. Figure 1

Steps in structure determination of tubular crystals grown from EDTA-treated YiiP. (A) Representative micrograph. (B) Selection of 2D class averages showing different helical symmetries: purple outline for tubes with D1 symmetry, green outline for tubes with D3 symmetry, and yellow outline for other, uncharacterized symmetries. (C) Helical reconstructions of tubular crystals with D1 and D3 symmetry. One dimer from the helical array has been colored. (D) Masked dimers from each of the reconstructions colored according to the local resolution ranging from 4.1 to 6.5 Å. The FSC plot for each reconstruction is also shown indicating an overall resolution of 4.2 Å.

### Suppl. Figure 2

Parametrization and validation of non-bonded dummy model for $Zn^{2+}$. (A) Geometry of the non-bonded dummy model. The central ZND site (magenta) is surrounded by six dummy sites DM (gray) forming a rigid octahedral shell. ZND-DM distances were maintained at 0.9 Å. Each DM particle carries a charge of $\delta^{+}$ = +0.35*e* and the central ZND particle carries –0.10*e* for a total charge of +2*e*. ZND and DM particles participate in van der Waals interactions. The approximate van der Waals radii of the ZND and DM particles are shown in the space-filling representation. (B) Results from the genetic algorithm parameter search. The Lennard-Jones parameters for the van der Waals interactions of the ZND and DM particles were optimized using a genetic algorithm to reproduce the $Zn^{2+}$ hydration free energy at standard conditions (HFE) of -1955 kJ/mol and the ion-oxygen distance (IOD) of 2.08 Å in simulations with water. The optimization ran through six iterations at which point it reproduced the experimental values within 1% and 0.3%, respectively. (C) Zn dummy model in water. The radial distribution function (RDF) for the Zn (ZND)-water oxygen distance, g(r), and its integral, N(r), show the first peak at the target IOD and the experimental number of water molecules in the first hydration shell, $n_1$ = 6. (D) Validation with MD simulations of $\beta$-1,3-1,4-endoglucanase (PDBID 1U0A, resolution 1.64 Å). The Zn binding site (with ZN 5005) is located in a dimer interface between chains A and C and formed by HHDD residues. (E) Binding site RMSD's of the three repeat simulations of 1U0A as time series and distribution over all repeat simulations (computed as a KDE): backbone of the protein in blue, backbone and sidechain atoms of the coordinating residues in orange, coordinating atoms only ($O_{\delta1}$/$O_{\delta2}$ for Asp, $N_{\varepsilon}$ or $N_{\delta}$ for His) in green. (F) Radial distribution of the distances of coordinating atoms (O and N) in 1U0A to the Zn ion (measured to the ZND center). Distributions were computed for each simulation repeat separately and averaged. The solid line shows the mean with bands, which are barely visible, indicating the standard deviation. The dashed vertical lines mark the distances in the crystal structure. (G) Validation with MD simulations of stromelysin-1 (PDBID 2USN, resolution 2.2 Å). The Zn binding site (with ZN 258) is formed by HHHD residues. (H) Binding site RMSD's of the three repeat simulations of 2USN as in E. (I) Radial distribution of the distances of coordinating O and N atoms in 2USN, as in F. (J) Validation with MD simulations of $\delta$' subunit of *E. coli* clamp-loader complex (PDBID 1A5T, resolution 2.2 Å). The Zn binding site (with ZN 501) is formed by CCCC residues. (K) Binding site RMSD's of the three repeat simulations of 1A5T as in E. The coordinating atoms were the $C_{\gamma}$ of the four Cys residues. (L) Radial distribution of the distances of coordinating S atoms in 1A5T, otherwise as in F.

### Suppl. Figure 3

Steps in structure determination of the untreated YiiP/Fab complex from the SP1 data set. (A) Representative micrograph. (B) Selection of 2D class averages showing multiple views of the complex. (C) Heterogeneous refinement job in cryoSPARC was used to look for multiple conformations and to select a homogeneous subset of particles for high resolution refinement. (D) Non-uniform refinement job in cryoSPARC colored according to local resolution, which ranged from 3.5 - 7.0 Å for the best class. FSC plot indicates an overall resolution of 3.8 Å when C2 symmetry was imposed.

### Suppl. Figure 4

Stability of Zn binding sites in MD simulations of YiiP. The RMSD of binding site residues, represented by their non-hydrogen atoms in the backbone and sidechain, was calculated after optimal superposition of each protomer for apo and holo simulations and each simulation repeat (md0 – md2), shown in shades of purple and cyan, respectively. To improve readability, the time series were averaged over blocks of 20 ns with the solid line showing the mean and the error band containing 95% of the data. RMSD distributions generated by a KDE are shown to the right of the time series. On the far right, each binding site in the SP3sym structure is shown with the Zn ion (magenta sphere) and the coordinating residues (licorice representation). (A) Site A in the TMD. (B) Site B in the M2/M3 loop of the TMD. (C) Site $C_1$ in the CTD. (D) Site $C_2$ in the CTD.

### Suppl. Figure 5

RMSD plot for the remaining two MD simulations (md0 and md2) of the holo state. The three traces correspond to the different alignment schemes relative to the starting model (5VRF) as described in the main figure.

### Suppl. Figure 6

Steps in structure determination of the EDTA-treated YiiP/Fab complex from the SP2 data set. (A) Representative micrograph. (B) Selection of 2D class averages showing multiple views of the complex. (C) Heterogeneous refinement job in cryoSPARC was used to look for multiple conformations and to select a homogeneous subset of particles for high resolution refinement. (D) Non-uniform refinement job in cryoSPARC colored according to local resolution, which ranged from 4.0 - 7.0 Å for the best class. FSC plot indicates an overall resolution of 4.0 Å, which was determined without imposition of symmetry.

### Suppl. Figure 7

RMSD plots for the remaining two MD simulations (md0 and md2) of the apo state. The three traces correspond to the different alignment schemes relative to the starting model (5VRF) as described in Fig. 3.

### Suppl. Figure 8

Quantitation of M2/M3 loop movements during the MD simulations showing greatly enhanced dynamics in the apo state. (A-C) $C_\alpha$ RMSD for each protomer from the individual simulations (md0 - md2) after alignment of the TMD relative to the starting structure (5VRF). (D) $C_\alpha$ RMSD distributions for loop motion relative to the TMD, representing all the simulations as calculated by KDE. (E-G) $C_\alpha$ RMSD for each protomer after alignment of the loop relative to the starting structure. (H) $C_\alpha$ RMSD distributions for the intrinsic loop motion, representing all the simulations in the apo and holo states (repeated from Fig. 5B to aid comparison).

### Suppl. Figure 9

Steps in structure determination of the EDTA-treated YiiP/Fab complex from the SP3 data set. (A) Representative micrograph. (B) Selection of 2D class averages showing multiple views of the complex. (C) Heterogeneous refinement job in cryoSPARC revealed the presence of two distinct conformations, which were individually used for high resolution refinement. (D) Non-uniform refinement jobs for each class in cryoSPARC colored according to local resolution, which ranged from 3.0 - 7.0 Å. FSC plots indicate an overall resolution of 3.4 and 4.2 Å for the symmetric conformation and the bent conformation, respectively. (E) Three principle components derived from 3D variability analysis of the combined particles from both classes.

### Suppl. Figure 10

Steps in structure determination of the YiiP/Fab complex treated with both EDTA and TPEN from the SP4 data set. (A) Representative micrographs from two different imaging sessions. (B) Heterogeneous refinement job in cryoSPARC revealed the presence of two distinct conformations, which were individually used for high resolution refinement. (C) Non-uniform refinement job in cryoSPARC colored according to local resolution, which ranged from 3.3 - 7.0 Å. FSC plots indicate an overall resolution of 3.5 and 4.7 Å for the symmetric conformation and the bent conformation, respectively. (E) Three principal components derived from 3D variability analysis of the combined particles from both classes.

### Suppl. Figure 11

Dynamics of the CTD are influenced by Zn binding. (A) Overlay of CTD's from the indicated cryo-EM structures showing close overlap of secondary structural elements from the symmetric, holo structures: SP1, SP3sym and SP4sym. The CTD from SP3bent also matches the configuration from the holo state, which correlates with the presence of density at Zn site C (c.f., Fig. 7g). (B) The CTD's from SP2 and SP4bent (apo) are farther apart (16 Å between $C_\alpha$ atoms of Arg237 and Glu281) than those from SP3sym (12.2 Å) and SP3bent (11.1 Å), suggesting that Zn binding at site C has a stabilizing influence on this domain. (C) The distance between $C_\alpha$ atoms from Arg237 and Glu281 was used as a collective variable to monitor CTD dynamics during the MD simulations. Data from all three simulations are plotted for apo and holo states with distributions from KDE calculations shown on the right. Timeseries were averaged over 20-ns intervals, with the solid line showing the average and the error bands contain 95% of the data. These data indicate that there are larger fluctuations in the CTD for the apo state**.**

### Suppl. Figure 12

Display of 3D variability in the SP4 data set. (A) Side view of 11 (from a total of 20) structures generated from one of the principal components from the 3D variability analysis. This series captures the conformational transition from the C2 symmetric conformation to the asymmetric conformation with a bent TM domain. The CTD's from each structure have been aligned. Helices in the TMD are rainbow colored from blue to red, the CTD is grey and the Fab chains are orange and red. (B) Front view of the same 11 structures showing the bending of the M2 helix (cyan indicated by arrow) during the conformational transition. (C) View from the cytoplasm toward Zn site A for a subset of 5 structures from the 3D variability analysis illustrating the bending of M2 (cyan) and M5 (orange) during the conformational transition. (D) Plots of distances between $C_\alpha$ atoms from Leu154 and Leu199 on the cytoplasmic side of the TMD and between Ala43 and Ala185 on the periplasmic side for each of the components derived from 3D variability analysis of the SP4 data set. These plots are analogous to those in Fig. 8. Evidence of TM gate closure can be seen in components 1 and 2.

### Movie Suppl. 1

Animation showing component 0 of the 3D variability analysis for the SP3 data set.

### Movie Suppl. 2

Animation showing component 1 of the 3D variability analysis for the SP3 data set.

### Movie Suppl. 3

Animation showing component 2 of the 3D variability analysis for the SP3 data set.

### Movie Suppl. 4

Animation showing component 0 of the 3D variability analysis for the SP4 data set.

### Movie Suppl. 5

Animation showing component 1 of the 3D variability analysis for the SP4 data set.

### Movie Suppl. 6

Animation showing component 2 of the 3D variability analysis for the SP4 data set.

### Movie Suppl. 7

Animation showing a morph of the atomic models produced by Namdinator for component 1 from the 3D variability analysis of the SP4 data set (c.f., Fig. 7 - Fig. Suppl. 4). These objectively built structures show the transition from the symmetric, holo state to the asymmetric apo state, including bending of the CTD relative to the TMD and bending of M2 and M5 helices.

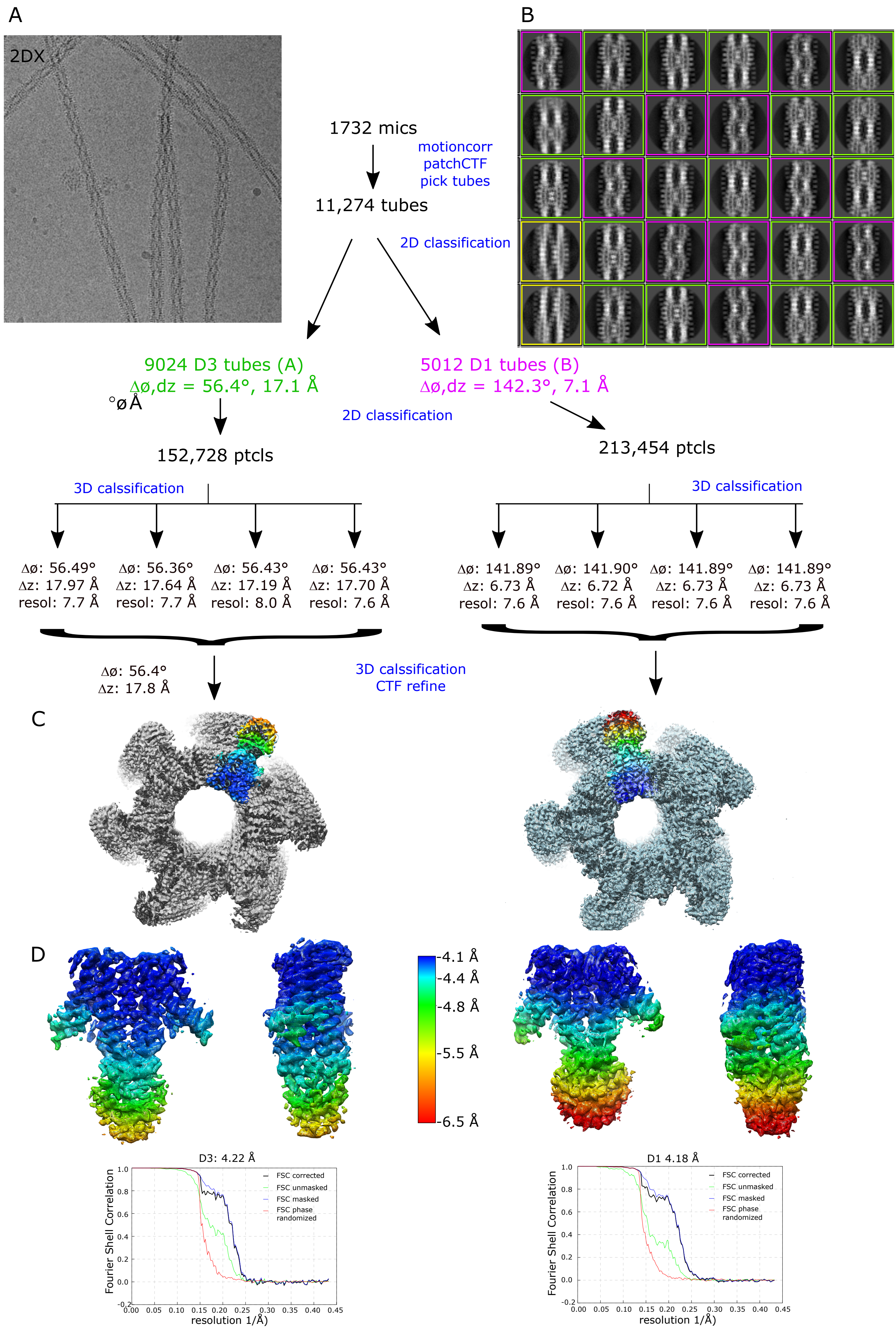

A
2DX
B
1732 mics
motioncorr
patchCTF
pick tubes
11,274 tubes
2D classification
9024 D3 tubes (A)
Δø,dz = 56.4°, 17.1 Å
°øÅ
5012 D1 tubes (B)
Δø,dz = 142.3°, 7.1 Å
2D classification
152,728 ptcls
213,454 ptcls
3D calssification
3D calssification
Δø: 56.49° Δz: 17.97 Å resol: 7.7 Å
Δø: 56.36° Δz: 17.64 Å resol: 7.7 Å
Δø: 56.43° Δz: 17.19 Å resol: 8.0 Å
Δø: 56.43° Δz: 17.70 Å resol: 7.6 Å
Δø: 141.89° Δz: 6.73 Å resol: 7.6 Å
Δø: 141.90° Δz: 6.72 Å resol: 7.6 Å
Δø: 141.89° Δz: 6.73 Å resol: 7.6 Å
Δø: 141.89° Δz: 6.73 Å resol: 7.6 Å
Δø: 56.4° Δz: 17.8 Å
3D calssification
CTF refine
C
D
-4.1 Å
-4.4 Å
-4.8 Å
-5.5 Å
-6.5 Å
D3: 4.22 Å
D1 4.18 Å
FSC corrected
FSC unmasked
FSC masked
FSC phase randomized
Fourier Shell Correlation
resolution 1/Å)


Suppl. Fig. 1

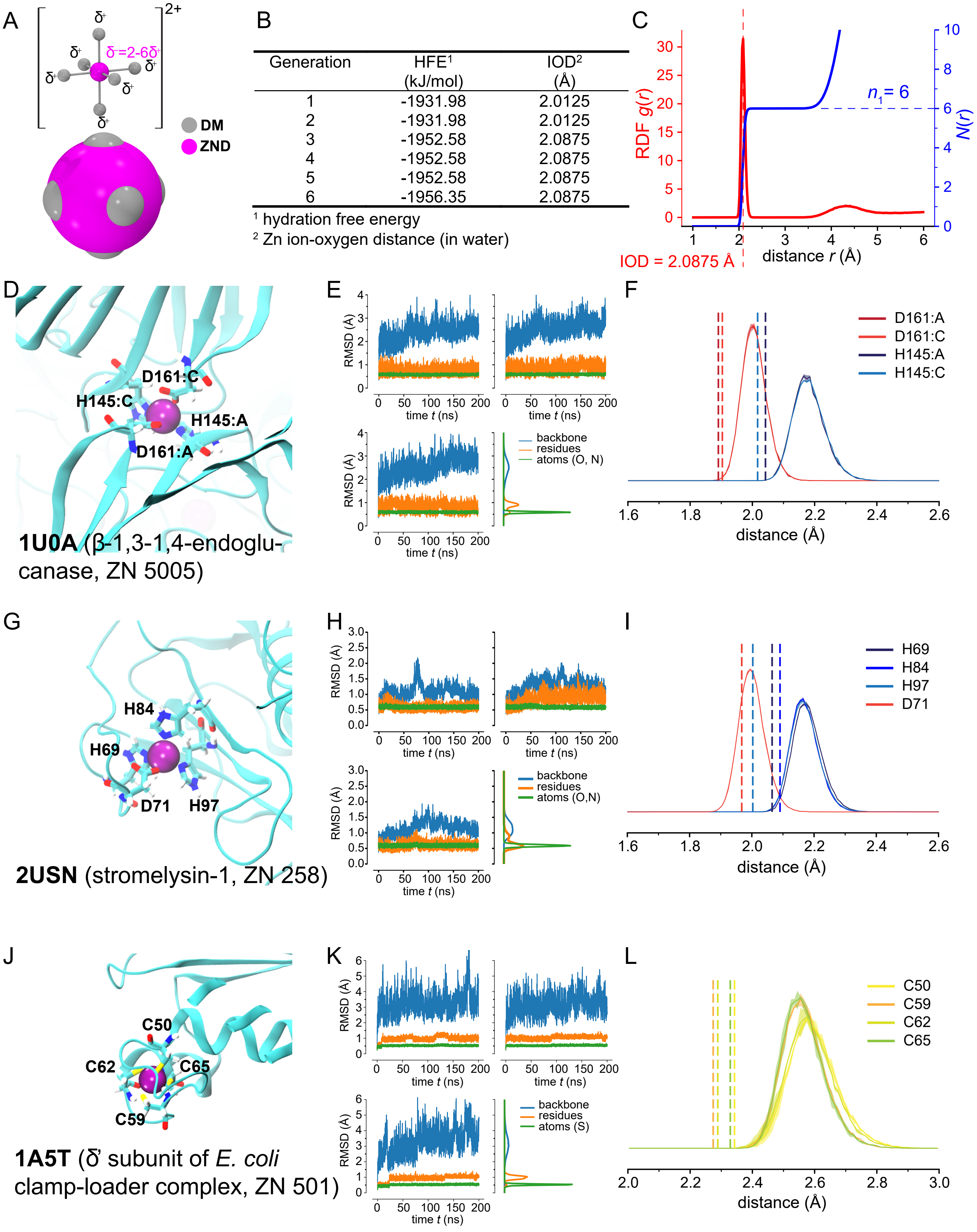


B

| Generation | HFE[1] (kJ/mol) | IOD[2] (Å) |
|---|---|---|
| 1 | -1931.98 | 2.0125 |
| 2 | -1931.98 | 2.0125 |
| 3 | -1952.58 | 2.0875 |
| 4 | -1952.58 | 2.0875 |
| 5 | -1952.58 | 2.0875 |
| 6 | -1956.35 | 2.0875 |

[1] hydration free energy
[2] Zn ion-oxygen distance (in water)

Suppl. Fig. 2

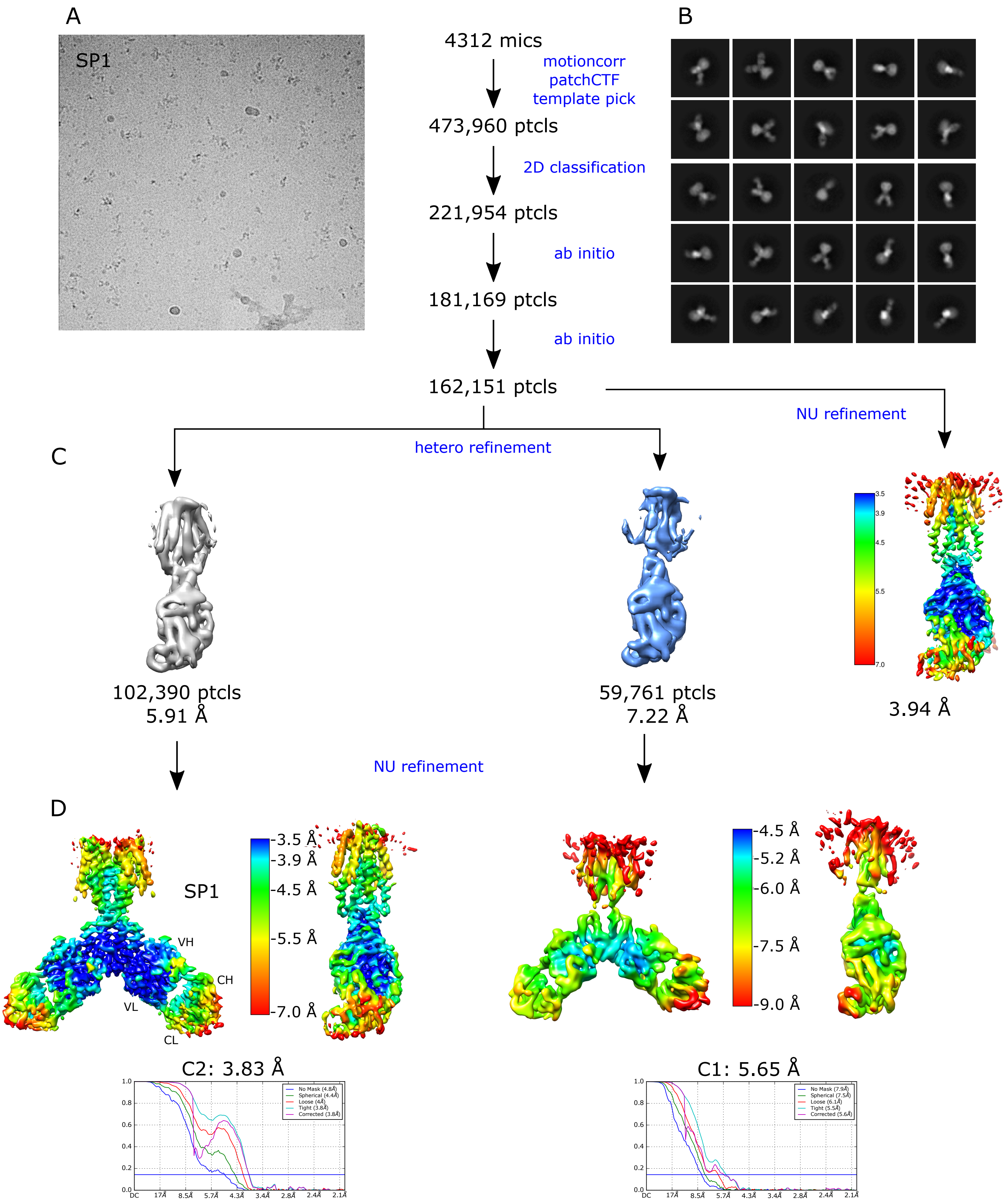
A
SP1
4312 mics
motioncorr
patchCTF
template pick
473,960 ptcls
2D classification
221,954 ptcls
ab initio
181,169 ptcls
ab initio
162,151 ptcls
B
NU refinement
C
hetero refinement
102,390 ptcls
5.91 Å
59,761 ptcls
7.22 Å
3.94 Å
NU refinement
D
SP1
VH
CH
VL
CL
3.5 Å
3.9 Å
4.5 Å
5.5 Å
7.0 Å
4.5 Å
5.2 Å
6.0 Å
7.5 Å
9.0 Å
C2: 3.83 Å
C1: 5.65 Å

Suppl. Fig. 3

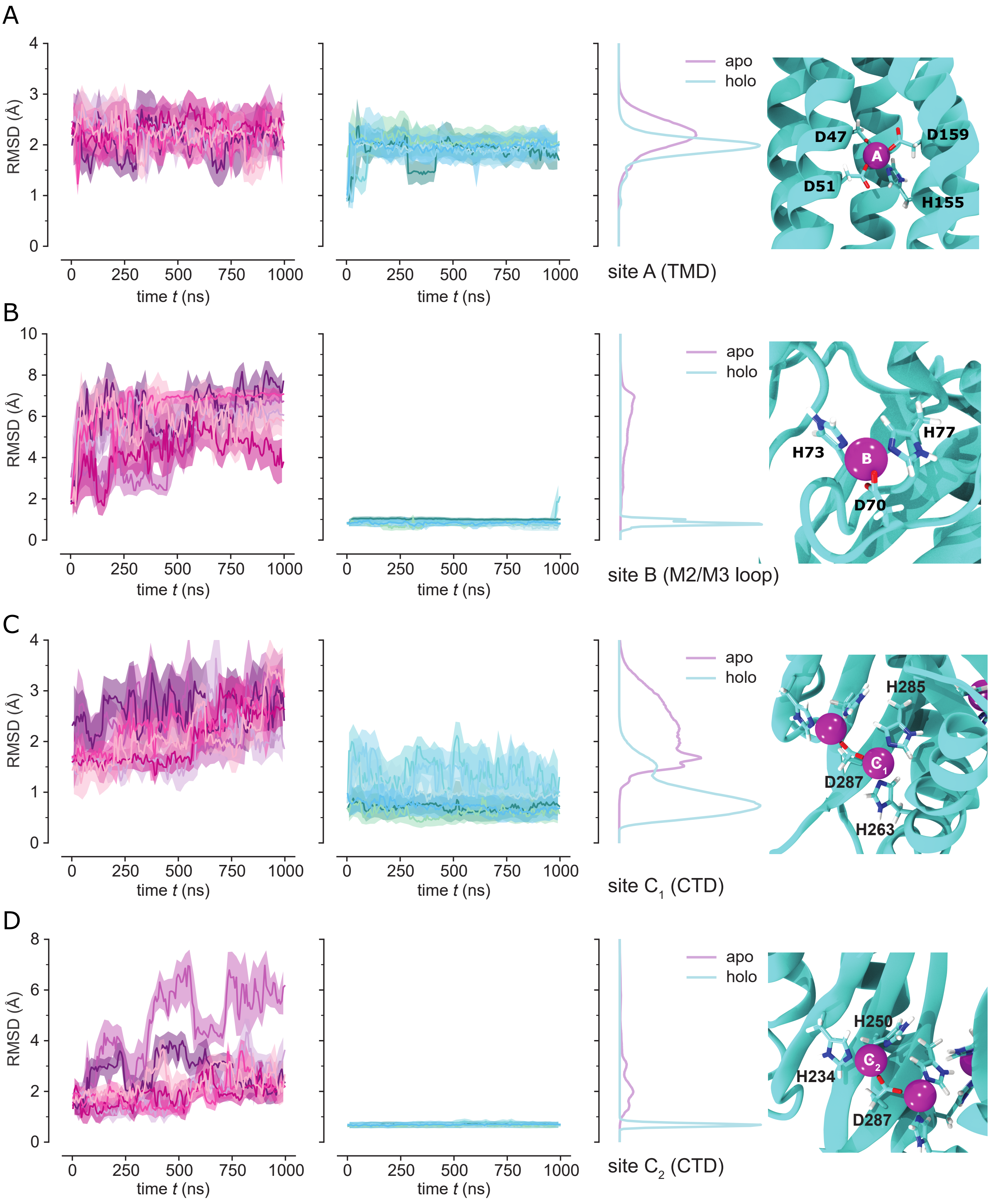
A
RMSD (Å)
time t (ns)
apo
holo
D47
D159
A
D51
H155
site A (TMD)
B
H73
H77
B
D70
site B (M2/M3 loop)
C
H285
$C_1$
D287
H263
site $C_1$ (CTD)
D
H250
$C_2$
H234
D287
site $C_2$ (CTD)

Suppl. Fig. 4

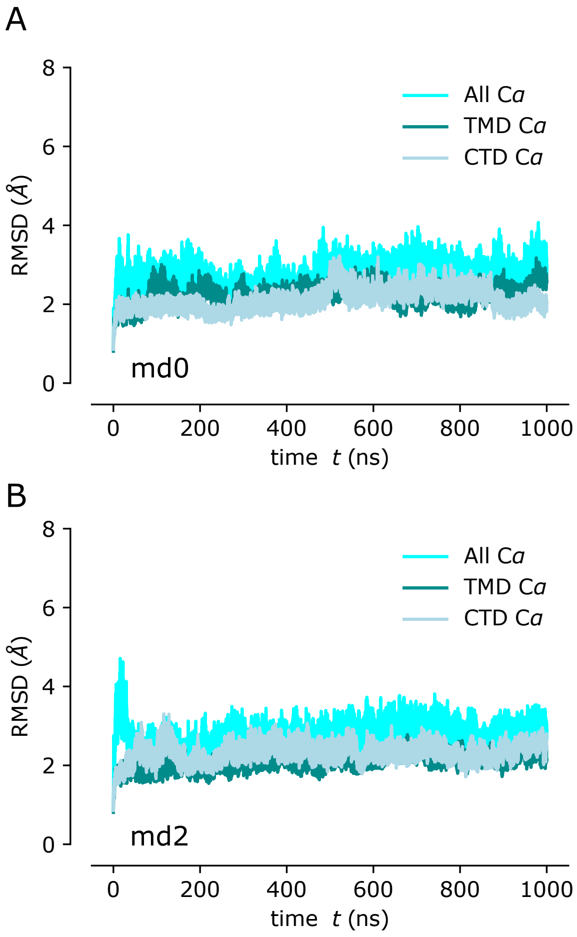
A
8
6
4
2
0
RMSD (Å)
All Cα
TMD Cα
CTD Cα
md0
0
200
400
600
800
1000
time t (ns)
B
8
6
4
2
0
RMSD (Å)
All Cα
TMD Cα
CTD Cα
md2
0
200
400
600
800
1000
time t (ns)

Suppl. Fig. 5

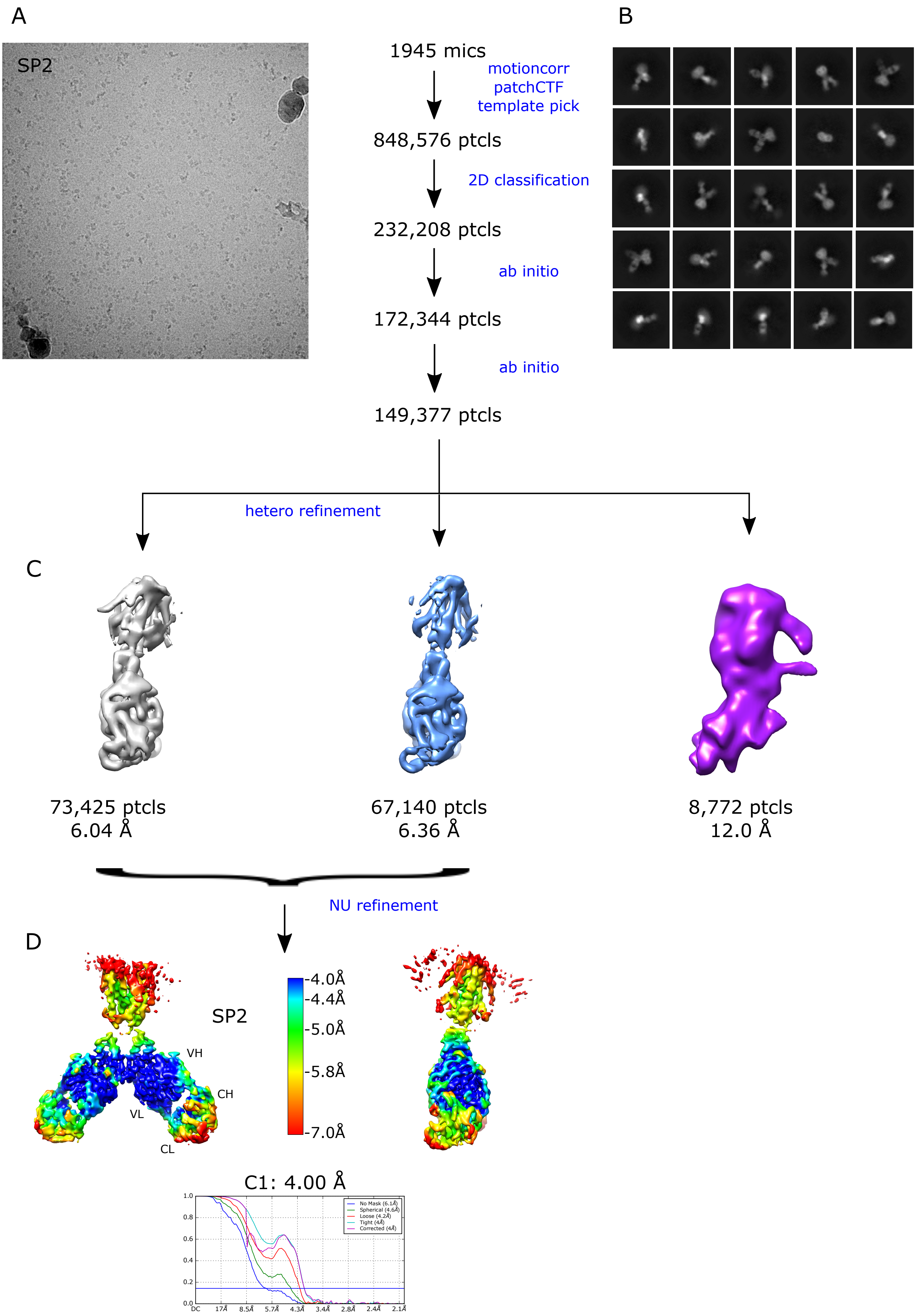
A
B
SP2
1945 mics
motioncorr
patchCTF
template pick
848,576 ptcls
2D classification
232,208 ptcls
ab initio
172,344 ptcls
ab initio
149,377 ptcls
hetero refinement
C
73,425 ptcls
6.04 Å
67,140 ptcls
6.36 Å
8,772 ptcls
12.0 Å
NU refinement
D
SP2
VH
CH
VL
CL
-4.0Å
-4.4Å
-5.0Å
-5.8Å
-7.0Å
C1: 4.00 Å

Suppl. Fig. 6

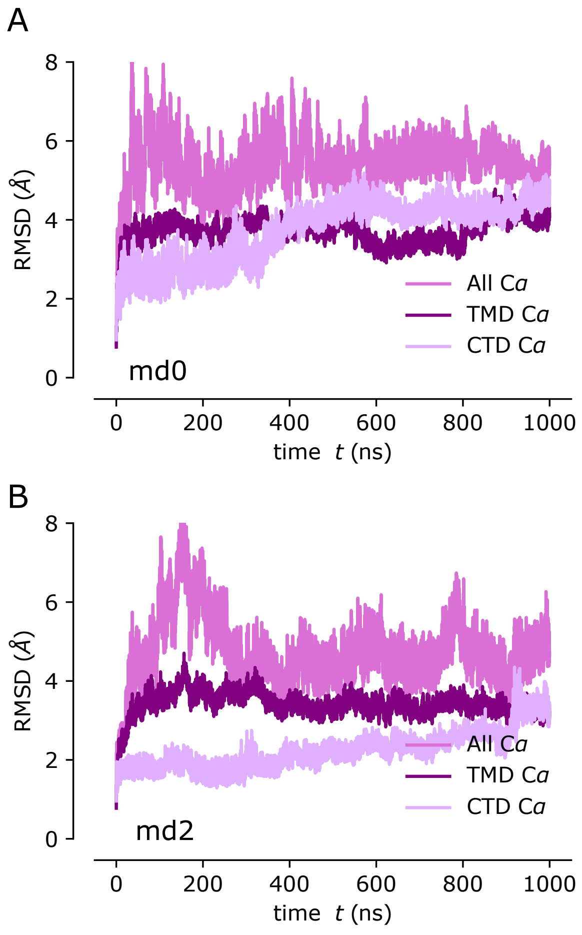
A
8
6
4
2
0
RMSD (Å)
All Cα
TMD Cα
CTD Cα
md0
0
200
400
600
800
1000
time t (ns)
B
8
6
4
2
0
RMSD (Å)
All Cα
TMD Cα
CTD Cα
md2
0
200
400
600
800
1000
time t (ns)

Suppl. Fig. 7

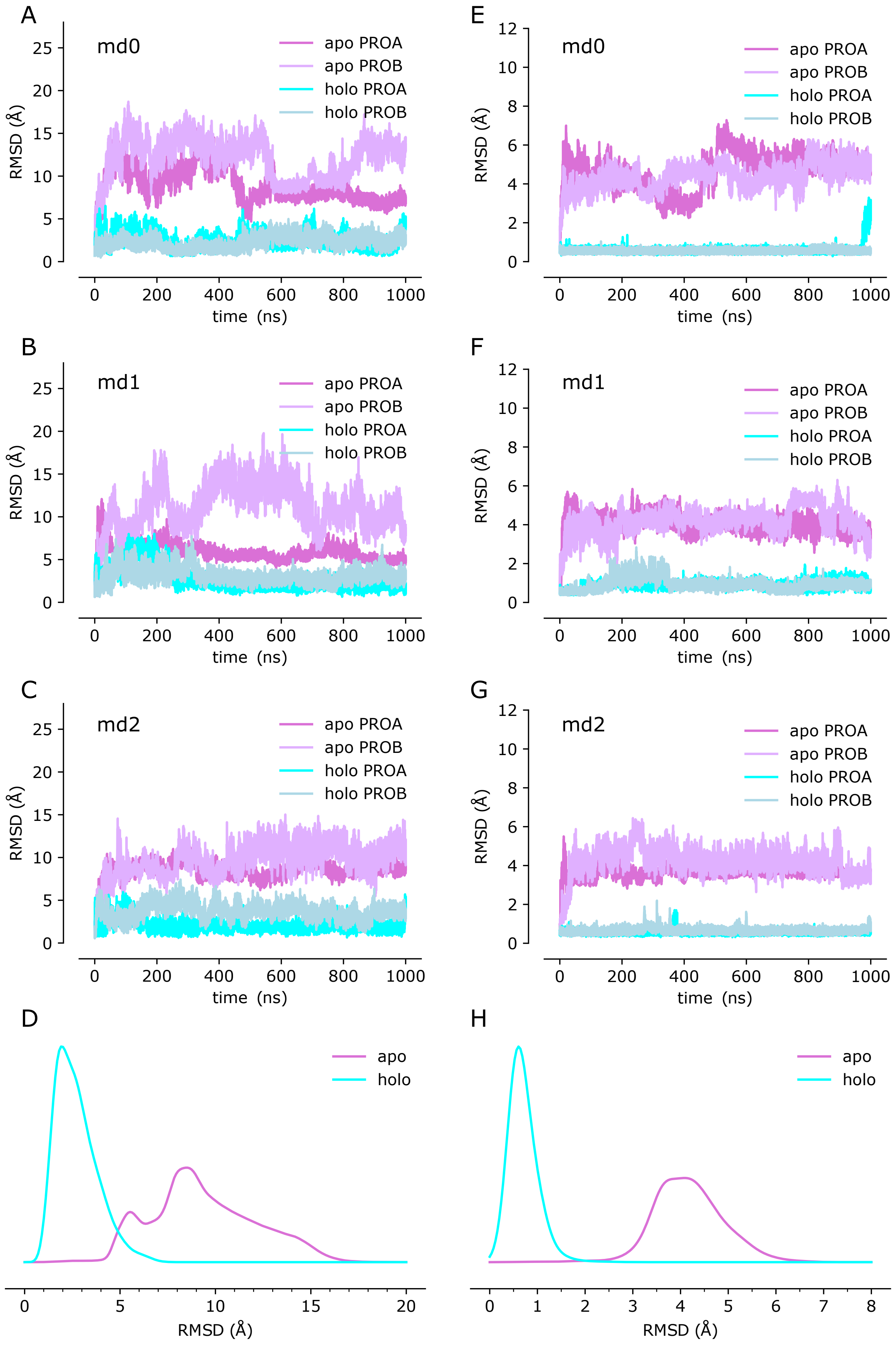
A
md0
apo PROA
apo PROB
holo PROA
holo PROB
RMSD (Å)
time (ns)
B
md1
C
md2
D
apo
holo
RMSD (Å)
E
F
G
H

Suppl. Fig. 8

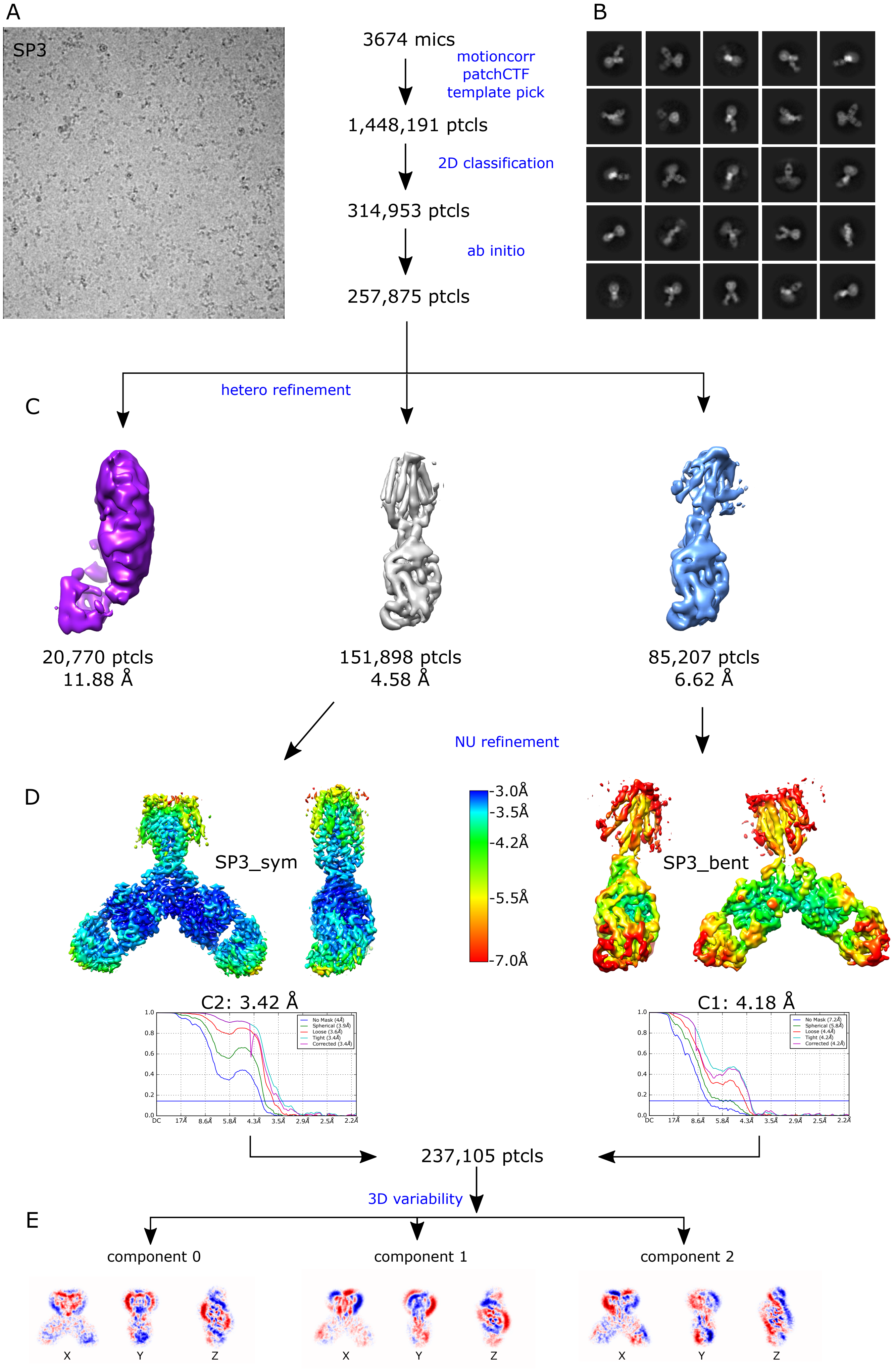
A
SP3
3674 mics
motioncorr
patchCTF
template pick
1,448,191 ptcls
2D classification
314,953 ptcls
ab initio
257,875 ptcls
B
hetero refinement
C
20,770 ptcls
11.88 Å
151,898 ptcls
4.58 Å
85,207 ptcls
6.62 Å
NU refinement
D
SP3_sym
-3.0Å
-3.5Å
-4.2Å
-5.5Å
-7.0Å
SP3_bent
C2: 3.42 Å
C1: 4.18 Å
237,105 ptcls
3D variability
E
component 0
component 1
component 2
X
Y
Z
X
Y
Z
X
Y
Z

Suppl. Fig. 9

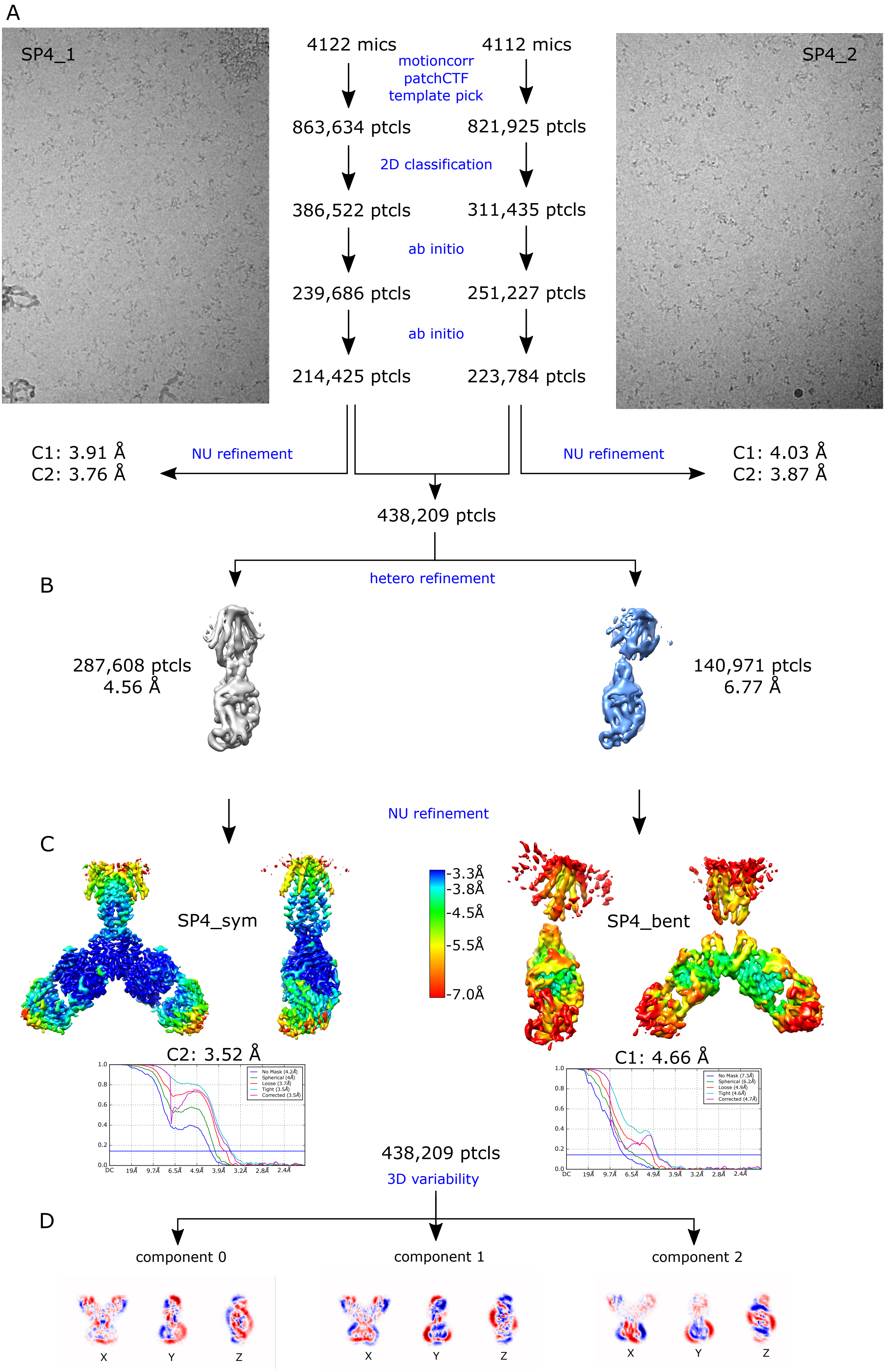

A
SP4_1
SP4_2
4122 mics
4112 mics
motioncorr
patchCTF
template pick
863,634 ptcls
821,925 ptcls
2D classification
386,522 ptcls
311,435 ptcls
ab initio
239,686 ptcls
251,227 ptcls
ab initio
214,425 ptcls
223,784 ptcls
C1: 3.91 Å
C2: 3.76 Å
NU refinement
NU refinement
C1: 4.03 Å
C2: 3.87 Å
438,209 ptcls
B
hetero refinement
287,608 ptcls
4.56 Å
140,971 ptcls
6.77 Å
NU refinement
C
SP4_sym
-3.3Å
-3.8Å
-4.5Å
-5.5Å
-7.0Å
SP4_bent
C2: 3.52 Å
C1: 4.66 Å
438,209 ptcls
3D variability
D
component 0
component 1
component 2
X
Y
Z


Suppl. Fig. 10

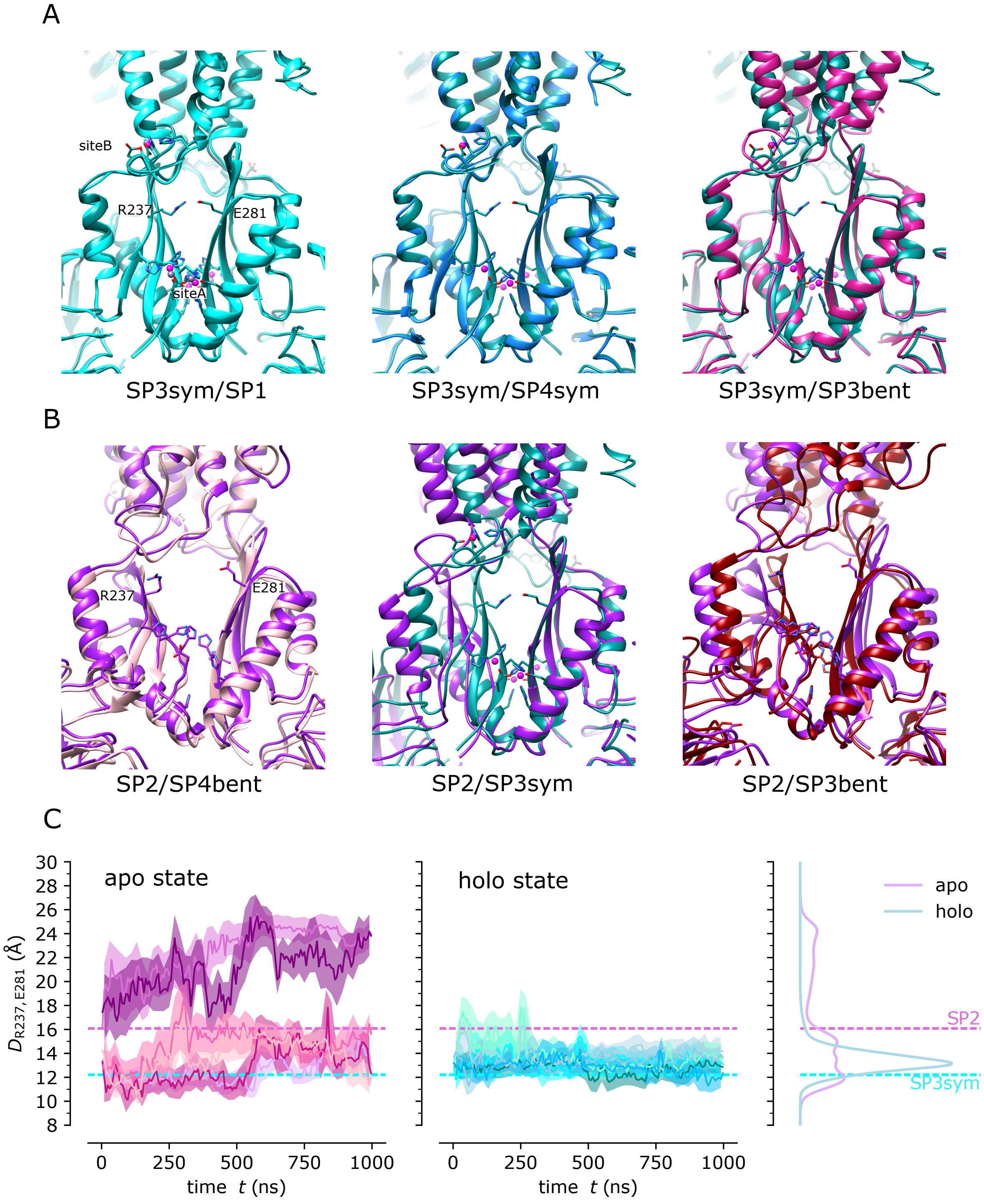
A
siteB
R237
E281
siteA
SP3sym/SP1
SP3sym/SP4sym
SP3sym/SP3bent
B
R237
E281
SP2/SP4bent
SP2/SP3sym
SP2/SP3bent
C
apo state
holo state
D R237, E281 (Å)
time t (ns)
time t (ns)
apo
holo
SP2
SP3sym

Suppl. Fig. 11

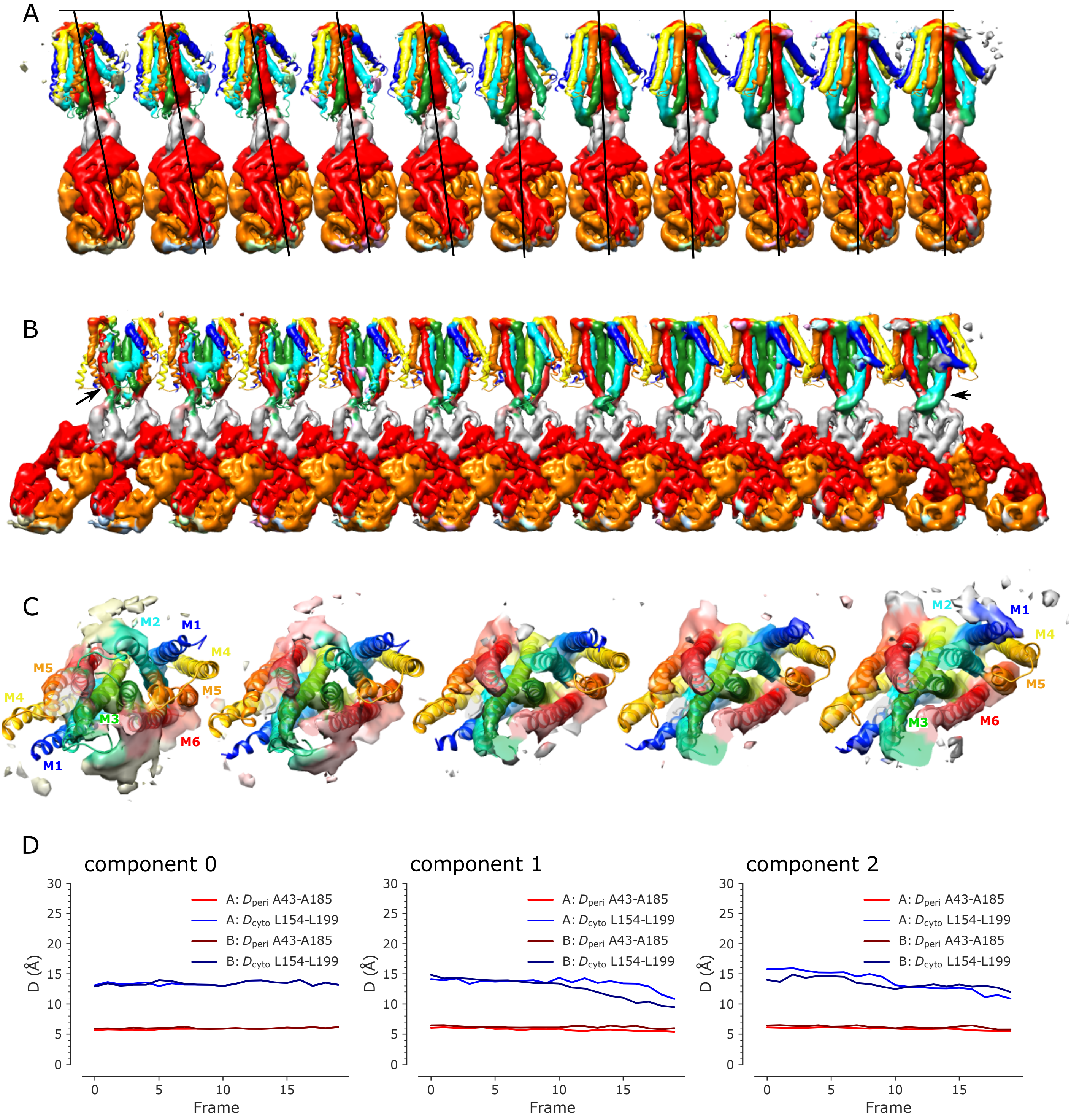
A
B
C
M2
M1
M4
M5
M4
M5
M3
M6
M1
M2
M1
M4
M5
M3
M6
D
component 0
component 1
component 2
A: $D_{peri}$ A43-A185
A: $D_{cyto}$ L154-L199
B: $D_{peri}$ A43-A185
B: $D_{cyto}$ L154-L199
D (Å)
Frame
0
5
10
15
20
25
30

Suppl. Fig. 12
